# Dendritic cell-Natural Killer cell Crosstalk Modulates T cell activation in Response to Influenza A Viral Infection

**DOI:** 10.1101/2022.08.12.503177

**Authors:** Abigail G. Harvey, Athens M. Graves, Chandana K. Uppalapati, Saoirse M. Matthews, Stephanie Rosenberg, Madison H. Fagerlie, Jack Guinan, Brina Lopez, Lisa M. Kronstad

## Abstract

Influenza viruses lead to substantial morbidity and mortality including ~3-5 million cases of severe illness and ~290,000-650,000 deaths annually. One of the major hurdles regarding influenza vaccine efficacy is generating a durable, robust cellular immune response. Appropriate stimulation of the innate immune system is key to generating cellular immunity. Crosstalk between innate dendritic cells (DC) and natural killer (NK) cells plays a key role in activating virus-specific T cells, yet the mechanisms used by influenza A viruses (IAV) to govern this process remain incompletely understood. Here, we used an *ex vivo* autologous human primary immune cell culture system to evaluate the impact of genetically distinct IAV strains on DC-NK cell crosstalk and subsequent T cell activation. We report that the addition of NK cells to cultures containing both DCs and naïve T cells led to an increase in the frequency of CD69^+^ and CD25^+^ T cells and elevated levels of IFN-γ, TNF, and IL-10. However, upon IAV infection of DCs, the addition of NK cells to cultures no longer increased the frequency of CD25^+^ T cells nor elevated IFN-γ, TNF, and IL-10 cytokine levels. Investigation of the impact of IAV infection on DC-NK crosstalk revealed that exposure of DCs to influenza virus in co-culture led to an increased frequency of HLA-DR^+^ and a decreased frequency of CD83^+^ and CD86^+^ cells–molecules involved in stimulating T cell activation. An expansion of an HLA-DR^+^ NK cell subset was observed following culture with influenza-infected DCs in a contact-dependent and cytokine independent-manner. Overall, our results indicate a role for DC-NK cell crosstalk in T cell priming in the context of influenza infection, informing the immunological mechanisms that could be manipulated for the next generation influenza vaccine or immunotherapeutic.

## Introduction

Influenza viruses cause annual seasonal epidemics and sporadic pandemics of zoonotic origin. The World Health Organization estimates the annual global burden of influenza at ~1 billion cases, including ~290,000-650,000 deaths. The U.S. economic burden of influenza-related illness has been estimated to reach $87 billion annually (1,2). The capacity of influenza viruses for rapid evolution via genetic reassortment and antigenic drift leads to evasion of immune surveillance by host antibodies specific to viral surface antigens. CD8^+^ T cells recognize viral peptides derived from internal viral components that are less subject to antibody-selected antigenic drift, and therefore more conserved across influenza strains and subtypes (3,4). Indeed, CD8^+^ T cells specific to conserved viral epitopes correlate with protection against symptomatic influenza illness, making a vaccine capable of stimulating heterologous CD8^+^ T cells an attractive target in vaccine design (5–9).

Initiating antigen-specific CD8^+^ T cell responses relies partly on dendritic cells (DC), which display cytosolic antigen on major histocompatibility (MHC) class I molecules via cross-presentation. Cross-presentation is crucial in activating effective adaptive immune responses to pathogens that do not typically infect DCs directly (10). DCs also shape the nature of the ensuing adaptive immune response depending on their specific phenotypic and functional state–inducing adaptive immune responses ranging from tolerogenic to stimulatory (11). The ability to engineer a desired adaptive immune response by exploiting the plasticity of DCs has made them a prime target in vaccine development. However, few studies have examined the possibility of DCs as an approach for universal influenza viral vaccine development, despite some promising studies demonstrating their success when used in populations most at risk for developing severe influenza-related illnesses such as immunocompromised individuals (12).

DC-based vaccines have been largely explored for cancer immunotherapy and less frequently, for the prevention of infectious diseases (13,14). While pre-clinical testing of DC vaccines has frequently shown promising results, clinical trials have faced challenges including induction of an effective activation state (15–20). Recent studies have highlighted the potential of employing natural killer (NK) cells to improve the potency of DC vaccine preparations (21,22). Bidirectional interactions between DCs and NK cells (crosstalk) influence both the magnitude and quality of the innate and adaptive immune responses (21,23,24). DCs act on NK cells to augment their cytotoxic potential, while NK cells, in turn, may act to counteract tumor or viral immune strategies by eliminating infected DCs with impaired antigen presentation functions. This may favor the development of DC-mediated CD4^+^ T_H_1 polarized response to antigens cross-presented by bystander DCs, which is responsible for protective immunity against intracellular pathogens (25,26). Indeed, NK cell depletion and reconstitution experiments showed that NK cell secretion of interferon (IFN)-γ was necessary for activation and expansion of T_H_1 CD4^+^ T cells following DC vaccination (27). In mice, Martín-Fontecha et al. showed that NK cells were rapidly recruited to secondary lymphoid organs–the location of DC-mediated naive T cell stimulation–after injection of the animals with lipopolysaccharide-matured DCs (27). This trafficking occurred in a manner dependent on the chemokine receptor CXCR3 (27). In humans, Draghi et al. found that influenza-infected DCs enhanced both NK cell expression of CD69, cytolytic activity, and IFN-γ production (28). Despite this knowledge, studies investigating the cellular changes that occur on T cells due to DC-NK cell crosstalk after influenza exposure are lacking.

In this study, we used an *ex vivo* autologous human triple cell co-culture system to characterize the phenotypic and functional profile of NK cells and DCs cultured alone and in concert, and to characterize their subsequent impact on T cell activation following exposure to either the 2009 H1N1 pandemic strain or the 2011 H3N2 seasonal strain. We found that NK cells and influenza A viruses (IAV) influence the expression of costimulatory (CD83 and CD86) and antigen presentation (HLA-DR) molecules on DCs and subsequently modulate the cytokine milieu and surface markers involved in T cell activation.

## Materials and Methods

### Virus Production and Titration

A/Victoria/361/2011 (Vic/11; H3N2) (BEI Resources) and A/California/07/2009 (Cal/09: H1N1) (BEI Resources) were propagated in 10-day-old specific pathogen-free embryonated chicken eggs (Charles River Laboratories Wilmington, MA) at 35°C and with 55-65% humidity. Allantoic fluid was harvested at 48 HPI followed by overnight incubation at 4°C. Each of the Vic/11 and Cal/09 batches was grown from a seed stock and multiple batches were tested. Infectious influenza titer was determined by a standard plaque assay on MDCK cells in the presence of 2 μg/mL L-1-tosylamide-2-phenylethyl chloromethyl ketone (TPCK)-treated trypsin. Influenza A virus inactivation was performed by delivering 2400 μJ/cm^2^ of UV light (254 nm) using a Strata linker 1800 (Stratagene, La Jolla, CA.). Virus inactivation was verified by plaque assay, or by intracellular influenza nucleoprotein antibody staining and analysis by flow cytometry.

### Peripheral Blood Mononuclear Cell Isolation

Whole blood (~350 mL) was collected from both male and female healthy adult donors via antecubital venipuncture into BD vacutainer, and Sodium Heparin blood collection tubes (BD, Franklin Lakes, NJ). Whole blood was diluted 2-fold in calcium- and magnesium-free phosphate-buffered saline (PBS pH: 7.2; Fisher Scientific, Waltham, MA) within 30 min of collection. A single-step density gradient centrifugation was performed to isolate peripheral blood mononuclear cells (PBMCs) by slowly layering diluted blood (30 mL) over 15 mL of Lymphoprep™ (STEMCELL Technologies, Vancouver, Canada) contained within either a 50 mL SepMate™ (STEMCELL Technologies) or a 50 mL conical tube. Stepwise centrifugation at 1,200 x *g* for 10 min at room temperature with slow deceleration was performed. The buffy coat interface containing PBMCs was collected into 50 mL conical tubes and pelleted via centrifugation (1,400 RPM for 10 min). Residual red blood cells were removed with treatment with ACK Lysing Buffer (Gibco, Waltham, MA). PBMCs were then washed twice with 1X PBS, pelleted as above, and cryopreserved at −80°C in 90% (v/v) fetal bovine serum (FBS; R&D Systems, Minneapolis, MN) and 10% (v/v) dimethyl sulfoxide (Corning, Corning, NY) in cryovials labeled with each donor’s unique donor sample identification number, followed by transfer to liquid nitrogen storage after 24 h.

### Differentiation of Monocytes into Monocyte-derived Dendritic cells (MoDCs)

Human MoDCs were generated using established protocols (29). Briefly, monocytes were purified from the PBMC population using Magnetic-Activated Cell Sorting (MACS^®^) with the Pan Monocyte Isolation kit (Miltenyi Biotech, Auburn, CA). Monocytes were then seeded at a density of 0.5 – 1.0 × 10^6^ cells/well. The monocyte population was further enriched by incubating cells in serum-free RPMI 1640 medium (Fisher Scientific, Waltham, MA) at 37°C and 5% CO_2_ for 2 h in 12-well Nunclon™ Delta surface-treated flat bottom plates (Thermo Fisher Scientific, Waltham, MA). Non-adherent cells were removed by gently washing the cells with pre-warmed culture medium. Adherent monocytes were then cultured with RPMI 1640 supplemented with 10% FBS, 1% L-Glutamine, and 1% Penicillin/Streptomycin (Thermo Fisher Scientific, Waltham, MA), recombinant human granulocyte-macrophage colony-stimulating factor, rhGM-CSF (800 U/L; Thermo Fisher Scientific, Waltham, MA) and recombinant human interleukin 4, rhIL-4 (500 U/L; Thermo Fisher Scientific, Waltham, MA) to induce MoDC differentiation. Media was changed on day two and day five of culture by removing the top half of the medium and replacing it with equal volumes of fresh RP10 medium supplemented with GM-CSF and IL-4.

### Cell Purification, Cell Stimulation, and IAV Infection

For each experiment, cryopreserved PBMCs were thawed and washed in RP10 medium (RPMI 1640 medium supplemented with 10% FBS plus 2 mM L-glutamine, 100 U/ml penicillin, and 100 U/ml streptomycin). Autologous naïve T cells and NK cells were purified by magnetic-activated cell sorting (MACS^®^; Miltenyi Biotec) according to the manufacturer’s instructions. Briefly, T cells were purified using the human Naïve Pan T Cell Isolation Kit (Miltenyi Biotec) and NK cells using the human NK Cell Isolation Kit (Miltenyi Biotec). The viability for each cell type was determined using a micro flow cytometer with propidium iodide (Moxi Flow; Orflo Technologies, Ketchum, ID) or a hemocytometer. MoDCs were then pelleted and re-suspended in serum-free RPMI, then plated into wells of a 96-well U-bottom tissue culture treated plate (CELLTREAT, Pepperell, MA) at a seeding density of 2 × 10^5^ cells/200uL/well and 0.5 × 10^5^cells/50uL/well. Cells were then infected with either Cal/09 or Vic/11 at a multiplicity of infection (MOI) of 3 for 1 h at 37°C and 5% CO_2_ in FBS-free RPMI. At 1 HPI, virus-containing supernatant was decanted, and MoDCs were either co-cultured with NK cells (1:4 or 4:1 ratios) and/or NK cells and pan naïve T cells (1:1:1, 4:1:1, or 1:4:1 ratios) in RP10 medium for the indicated time. NK cells or T cells stimulated with 162 nM Phorbol-Myristate-Acetate and 2.68 μM ionomycin (PMA/I) or MoDCs stimulated with 30 μg/mL Poly (I:C) (Invivogen) were used as positive controls. In transwell experiments, MoDCs and NK cells were seeded in a 24-well plate and separated during culture by a semipermeable membrane of 0.4-μm pore size (Corning).

### Flow Cytometry

Cell purity after magnetic isolation was accessed using indicated lineage markers and analytical flow cytometry. For MoDCs and NK cell co-culture experiments, cells were first stained with LIVE/DEAD™ Fixable Violet Dead Cell Stain kit (Thermo Fisher Scientific), followed by surface staining with APC anti-human CD7 (clone: CD7-6B7; BioLegend^®^), Phycoerythrin (PE)-Cy7 anti-human CD56 (clone: B159; BD Biosciences), PE anti-human CD83 (clone: HB15e; BioLegend^®^, San Diego, CA), Brilliant Violet 510™ anti-human CD86 (clone: IT2.2; BioLegend^®^), Brilliant Violet 570™ anti-human HLA-DR (clone: L243; BioLegend^®^), and PE anti-human HLA-A, B, C (clone: W6/32; BioLegend^®^) antibodies. For MoDC, NK cell, and T cell co-culture experiments, cells were first stained with Live-or-Dye™ fixable AmCyan (Biotium, Fremont, CA) followed by staining with PE anti-human CD3-phycoerythrin (clone: UCHT1; BioLegend^®^), Pacific Blue™ anti-human CD69 (clone: FN50; BioLegend^®^), and anti-CD25-Alexa-647 (clone: BC96; BioLegend^®^) antibodies. Fixation and permeabilization with FACS Lyse and FACS Perm II (BD Pharmingen) were performed according to the manufacturer’s instructions. Cells were stained with FITC anti-IAV Nucleoprotein antibody (Life Technologies, Waltham, MA) to determine the percentage of virally infected MoDCs. Uncompensated data were collected by analytical flow cytometry (Guava^®^ easyCyte™ 12HT flow cytometer (Luminex Corporation, Austin, TX)). Flow cytometry analysis and compensation were performed using FlowJo™ version 10.6.1 (BD Biosciences) software. Fluorescence Minus One (FMO) controls were included in each biological replicate where samples were stained with all fluorophores in the panel, except one. This corresponding sample was used to set the upper boundary for the background signal, permitting the gating of the positive population.

### RNA isolation and quantitative RT-PCR

NK cells cultured in isolation or co-cultured for 23 h with mock-treated, Cal/09- or Vic/11-infected MoDC at a 4:1 (MoDC+NK cell) ratio were purified by magnetic-activated cell sorting (MACS^®^) according to the manufacturer’s instructions (NK cell Isolation kit; Miltenyi Biotec). Total RNA from the samples was isolated using TRIzol^®^ reagent following the manufacturer’s isolation protocol (Life Technologies), by adding 5 μg of RNase-free glycogen as a carrier when precipitating RNA. RNA purity was verified by NanoDrop (Thermo Fisher Scientific) before DNase treatment and reverse transcription using SuperScript**™** IV VILO**™** Master Mix with ezDNase™ Enzyme (Invitrogen, Waltham, MA). Primer sequences for HLA-DRA (Forward *5’-AGG CCG AG TCT ATC TGA ATC CT-3’;* Reverse *5’-CGC CAG ACC GTC TCC TTC T-3’*) (30) and Cyclophilin (Forward *5’-TGC CAT CGC CAA GGA GTA G-3’;* Reverse *5’-TGC ACA GAC GGT CAC TCA AA-3’*) were used and HLA-DRA mRNA levels were normalized to Cyclophilin. Samples were analyzed on Bio-Rad CFX96 Touch Real-Time PCR Detection System (Bio-Rad, Hercules, CA) using Power Track SYBR Green master mix per manufacturer instructions (Thermo Fisher Scientific). Relative abundance of HLA-DRA transcript within samples was determined with the 2^-ΔΔCt^ method.

### Multiplex Human Cytokine Magnetic Bead Panel

Supernatants from three donors were collected at 96 HPI and were stored at −80°C. Samples were analyzed using a custom, commercially available multiplex magnetic assay kit per manufacturer instructions (Milliplex^®^; MilliporeSigma, Burlington, MA). Briefly, supernatants harvested from co-cultures (i.e., 1) T cells only; 2) MoDCs and T cells; 3) NK cells and T cells and; 4) MoDCs, NK cells, and T cells with ratios of 0:0:1, 1:0:1, 0:4:1, and 1:4:1) were transferred, in duplicates, to the provided 96-well plates to analyze cytokine concentrations. Each well received 25 μL assay buffer, followed by 25 μL of sample supernatant, and 60 μL magnetic beads. Background wells received 25 μL of assay buffer instead of 25 μL sample supernatant. Quality controls and high sensitivity human cytokine standards 1-7 were separately resuspended, inverted, and vortexed in 250 μL deionized water followed by serial dilution. The plate was sealed and incubated with shaking overnight at 4°C. After a 16 hr incubation period, the plate underwent triple washes with wash buffer before adding 25 μL of detection antibodies, incubated for 1 hr at 27°C, followed by adding 25 μL Streptavidin-Phycoerythrin to each well with detection antibodies. The plate was incubated for 30 min at 27°C and then washed three times. 150 μL of drive fluid was added to all wells. Magnetic beads were resuspended by using a plate shaker for five minutes. Cytokines of interest include: IFN-α2, IFN-γ, IL-2, IL-10, IL-12p70, IL-15, IL-18, TNF, and IL-21. Plates were analyzed on the MAGPIX instrument (Luminex) per manufacturer instructions. Minimum detectable concentrations for each cytokine were: IFN-α2, 2.9 pg/mL; IFN-γ, 0.8 pg/mL; IL-2, 1.0 pg/mL; IL-10, 1.1 pg/mL; IL-12p70, 0.6 pg/mL; IL-15, 1.2 pg/mL; IL-18, 0.94 pg/mL; TNF, 0.7 pg/mL; IL-21, 18.49 pg/mL. Concentrations were derived from standard curves set to a 5-parameter (log scale) curve fit.

### Statistical Analysis

A Tukey’s multiple comparison test on a two-way ANOVA using GraphPad Prism V9.0.0 was performed to analyze data for transwell, quantitative RT-PCR, and multiplex cytokine assays. Paired Wilcoxon signed-rank tests were performed using R. An alpha value of 0.05 was set to all data analysis and a p-value of < 0.05 was considered statistically significant.

### IRB Approval-Ethics

Blood from healthy adult donors, aged 18-65 years of age, both males and females, was used for these experiments. The research protocol for this study was approved by Midwestern University’s Institutional Review Board under study number IRB/AZ 1299. Additional PBMCs were obtained from leukoreduction system chambers purchased from Vitalant (Scottsdale, AZ) and STEMCELL Technologies. Because these subjects were fully de-identified, the study protocol was deemed not to fall under human subjects research by the Institutional Review Board at Midwestern University.

## Results

### Establishment of an autologous culture system to investigate monocyte-derived DC (MoDC)-NK cell crosstalk in response to IAV

We first established an autologous co-culture system to investigate human DC-NK cell crosstalk in response to A/California/07/2009 (H1N1) (Cal/09) and A/Victoria/361/2011 (H3N2) (Vic/11) IAV. These viruses were selected based on a prior report detailing that NK cells produced higher levels of IFN-γ in response to monocytes infected with Cal/09 compared to Vic/11, potentially leading to distinct outcomes during DC-NK cell crosstalk (31). To generate MoDCs, monocytes were purified from cryopreserved PBMCs using negative selection followed by culturing in the presence of IL-4 and GM-CSF. After seven days, the MoDCs displayed characteristic DC morphology under the light microscope, consisting of cytoplasmic projections and the formation of loosely adherent aggregates. Expression of the monocyte marker CD14 was decreased while expression of the pathogen recognition receptor CD209 was increased when compared to freshly isolated monocytes from the same representative donor–a surface marker expression pattern consistent with MoDCs **(Figure S1)**. Phenotypic signatures of MoDCs (CD7^-^) and NK cells (CD7^+^CD56^+^) were assessed by flow cytometry permitting lineage gating to allow for expression patterns on MoDCs and NK cells to be assessed individually. A representative flow cytometry gating schematic of MoDCs and NK cells is shown in **Figure 1**. MoDCs were then either mock-treated (uninfected) or infected with Cal/09 or Vic/11 IAV strains at an MOI of 3 followed by co-culture for 23 hours (h) with autologous NK cells. Infection levels in MoDCs were tracked by staining for intracellular expression of the influenza A nucleoprotein (FluA-NP). The frequency of FluA-NP^+^ MoDCs increased following exposure to both Cal/09 and Vic/11 at 24 h post-infection although not when the virus was inactivated with ultraviolet (UV) light **(Figure 1)**.

**Figure 1.**
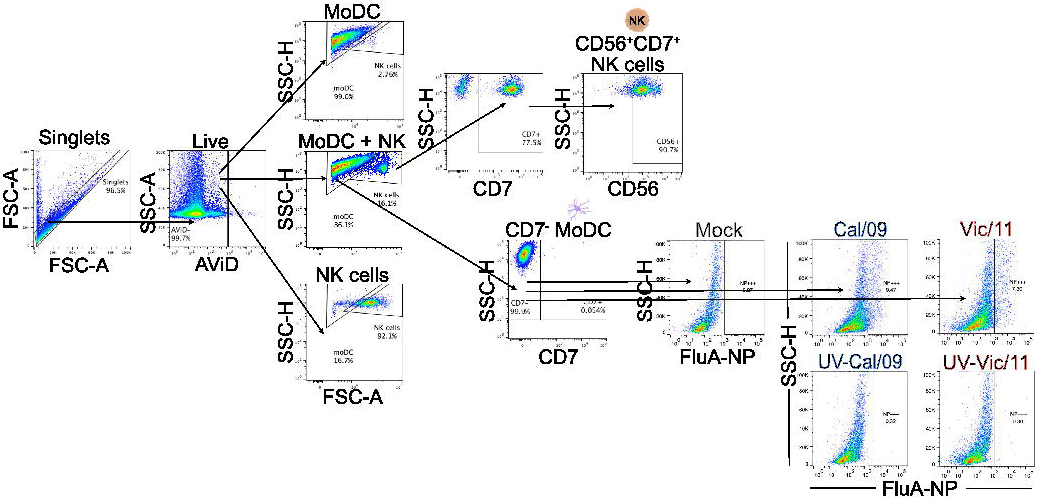
Lineage gating schematic for MoDCs and NK cells. Diagram of lineage gating tree used to identify MoDCs and NK cells by flow cytometry. Cal/09 and Vic/11 infection of MoDCs at an MOI of 3 at 24 HPI is shown using an antibody specific to IAV nucleoprotein (FluA-NP). AViD, LIVE/DEAD Fixable Violet Dead Cell Stain; FSC, forward scatter; SSC, side scatter.

### IAV decrease the expression of CD83 and CD86 and increase HLA-DR on MoDCs when co-cultured with NK cells

Using the MoDC-NK cell co-culture model established above, we first investigated whether the expression of two MoDC maturation-associated phenotypic markers important for activation of naïve T cells, CD83 and CD86, was influenced by exposure to either IAV strain, and whether expression was modified in the presence of NK cells. Polyinosinic-polycytidylic acid (Poly (I:C)), a double-stranded RNA mimic recognized by Toll-like receptor 3 (TLR3) and expressed by DCs, was used as a positive control to stimulate the MoDCs. As expected, Poly (I:C) treatment led to an increase in the frequency of CD83^+^ MoDCs, as compared to mock treatment **(Figure 2A-B)**. The outcome of DC-NK cell crosstalk has been reported to vary depending on cell ratio, with high DC to NK cell ratios amplifying DC cytokine production and low DC to NK cell ratios inhibiting DC function due to killing by autologous NK cells (editing) (32,33). For this reason, the impact of NK cells on the expression of DC surface markers was evaluated at two different DC to NK cell ratios– 1:4 or 4:1. In the presence of NK cells at either cell ratio, the frequency of CD83^+^ and CD86^+^ MoDCs was not significantly altered compared to when MoDCs were cultured alone **(Figure 2A-D)**. We next evaluated whether IAV exposure affected the frequency of CD83^+^ **(Figure 2A-B)** or CD86^+^ **(Figure 2C-D)** on MoDCs using the same design as above. At 24 h post-infection (HPI), exposure to either Cal/09 or Vic/11 led to a decrease in the percentage of MoDCs expressing CD83 at the higher MoDC to NK cell ratio (4:1), although not when cultured alone (1:0) or at the low MoDC to NK cell ratio (1:4) **(Figure 2A-B)**. Similar results were observed for the percentage of MoDCs expressing CD86 but only when exposed to Cal/09 **(Figure 2C-D)**. Statistically significant decreases in the percentages of MoDCs expressing CD83 and CD86 were not found at 7 HPI.

**Figure 2.**
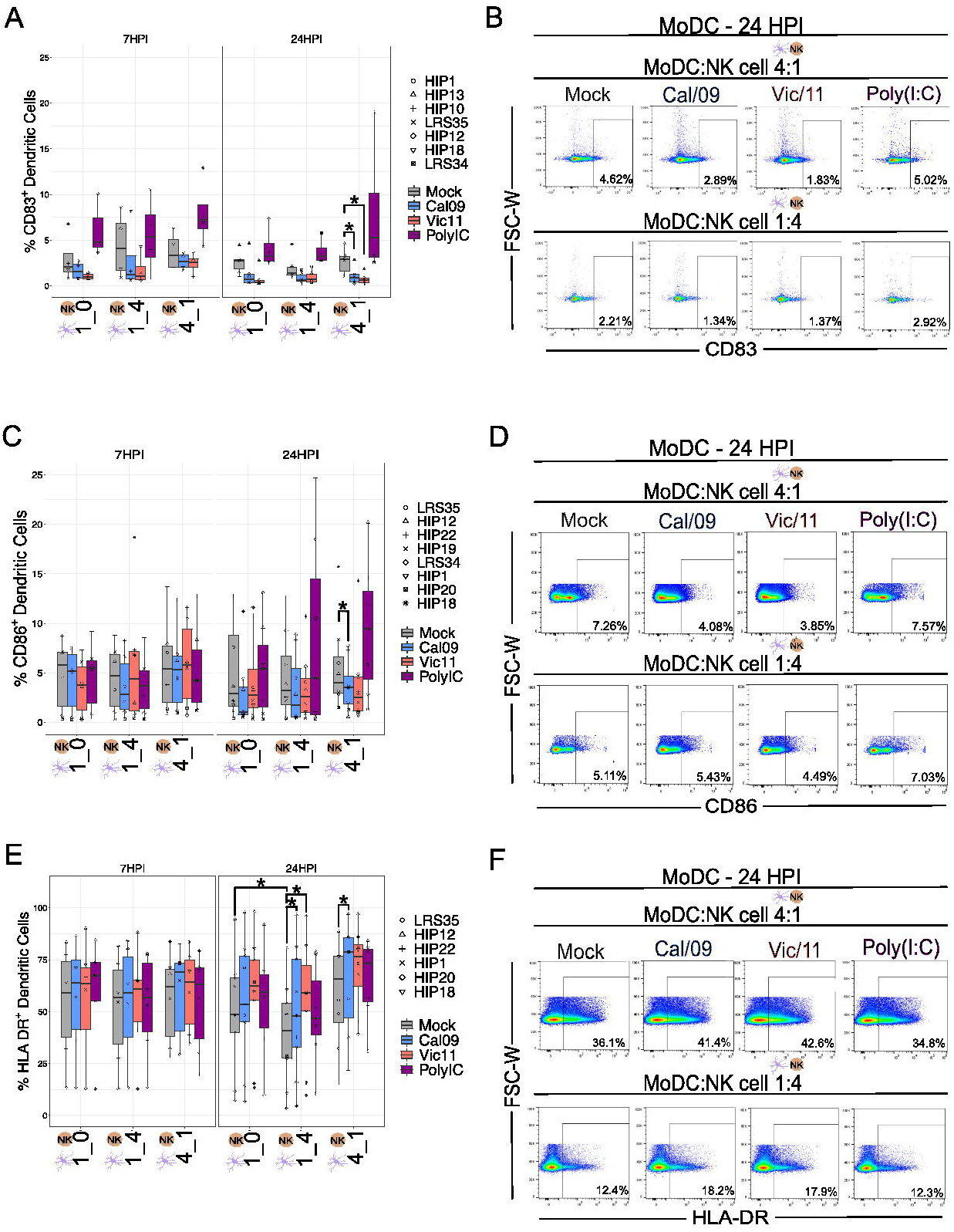
Downregulation of CD83 and CD86 costimulatory molecules and augmented antigen presentation by IAV-infected MoDCs in co-culture with NK cells. **(A)** Summary plot of Cal/09 or Vic/11-infected MoDCs (MOI = 3) CD83^+^ frequency after 24 HPI cultured alone (1:0) or after 7 or 23 h co-culture with NK cells at indicated cell ratio (*n* = 7) as assessed by flow cytometry using an antibody specific to CD83. **(B)** Representative flow plot of CD83 expression on MoDCs after 23 h co-culture with Cal/09- or Vic/11-infected MoDC (MOI = 3) at a MoDC to NK cell ratio of 4:1 (top panel) or 1:4 (bottom panel). **(C)** Summary plot of Cal/09- or Vic/11-infected MoDCs (MOI = 3) CD86^+^ frequency after 24 HPI cultured alone (1:0) after 7 or 23 h co-culture with NK cells (*n* = 8) as assessed by flow cytometry using an antibody specific to CD86. **(D)** Representative flow plot of CD86 expression on MoDCs after 23 h co-culture with Cal/09- or Vic/11-infected MoDCs (MOI = 3) at a MoDC to NK cell ratio of 4:1 (top panel) or 1:4 (bottom panel). **(E)** Summary plot of Cal/09 or Vic/11-infected MoDCs (MOI = 3) HLA-DR^+^ frequency after 24 HPI cultured alone or 7 or 23 h co-culture with NK cells (*n* = 6) as assessed by flow cytometry using an antibody specific to HLA-DR. **(F)** Representative flow plot of HLA-DR expression on MoDCs after 23 h co-culture with Cal/09- or Vic-11 infected MoDCs (MOI = 3) at a MoDC to NK cell ratio of 4:1 (top panel) or 1:4 (bottom panel). Poly (I:C) treatment served as a positive control. \**p* < 0.05, Wilcoxon signed-rank test.

We next investigated the impact of NK cells and IAV exposure on the MoDC expression of human leukocyte antigen (HLA)-DR–a subtype of HLA class II necessary for antigen presentation. Relocalization of HLA class II molecules occurs from late endosomal compartments to the plasma membrane. This process is increased when DCs acquire immunostimulatory properties such as expression of CD83 (34). HLA-DR expression was measured on MoDCs at either 7 or 24 HPI either in isolation or when in co-culture with NK cells at low (1:4) or high (4:1) MoDC to NK cell ratios. At 24 HPI, although not 7 HPI, the presence of a low MoDC to NK cell ratio (1:4) led to a decrease in the frequency of HLA-DR^+^ MoDCs compared to MoDCs cultured alone. In contrast, infection with Cal/09 at both cell ratios and with Vic/11 at the low MoDC to NK cell ratio (1:4) increased the percentage of HLA-DR^+^ MoDCs, although not when MoDCs were cultured alone (1:0) **(Figure 2E-F)**. No significant changes in the frequency of HLA-DR^+^ MoDCs were observed after Poly (I:C) treatment, consistent with results observed in prior reports (35–37). Under steady state, uninfected conditions, the percentage of HLA-DR^+^ MoDCs was decreased at low MoDC to NK cell ratios (1:4), although not at high MoDC to NK cell ratios (4:1) compared to when MoDCs were cultured in isolation. In contrast, IAV exposure led to an increase in the frequency of HLA-DR^+^ MoDCs at both cell ratios compared to uninfected MoDCs. Consistent with prior findings that IAV infection fails to downregulate HLA class I molecules, expression of HLA class I molecules appeared similar on IAV-infected MoDCs compared to mock treatment **(Figure S2)** (38). Taken together, these data showed that while exposure of MoDCs to IAV increased HLA-DR, in contrast, CD83 and CD86 maturation markers were reduced, consistent with a tolerogenic-like DC phenotype (39).

### Influenza A virus-infected MoDCs induce HLA-DR expression on NK cells

Expression of HLA-DR is generally restricted to professional antigen-presenting cells–DCs, macrophages, and B cells–although its expression has been documented on NK cells (40). For example, an HLA-DR^+^ NK cell subset was found to be expanded in the peripheral blood of patients with primary tuberculosis and to mediate cytokine production by CD4^+^ T cells (41). We therefore investigated HLA-DR expression on either NK cells alone or in co-culture with MoDCs at 7 or 24 HPI. Phorbol-Myristate-Acetate and Ionomycin (PMA/I) was used as a positive control for chemical stimulation. At 7 and 24 HPI, there was an increase in the frequency of HLA-DR^+^ NK cells when cultured with uninfected MoDCs, with markedly higher HLA-DR expression observed at the high MoDC to NK cell ratio (4:1) and to a lesser extent at the low MoDC to NK cell ratio (1:4) **(Figure 3A-B)**. Upon MoDC infection with IAV, the percentage of HLA-DR^+^ NK cells increased further, independent of the IAV strain used to infect MoDCs (**Figure 3A-B)**. Exposure to influenza virions was insufficient to change the frequency of NK cells expressing HLA-DR in the absence of MoDCs (MoDC to NK cell ratio of 0:1) **(Figure 3A-B**). In contrast to the increase in HLA-DR expression, NK cells failed to acquire expression of either CD83 **(Figure S3A-B)** or CD86 **(Figure S3C-D)** costimulatory molecules after exposure to uninfected or to IAV-infected MoDCs. Collectively, the culture of NK cells with MoDCs leads to an increase in HLA-DR expression by NK cells, which is markedly increased when the MoDCs have been infected with either IAV strain.

**Figure 3.**
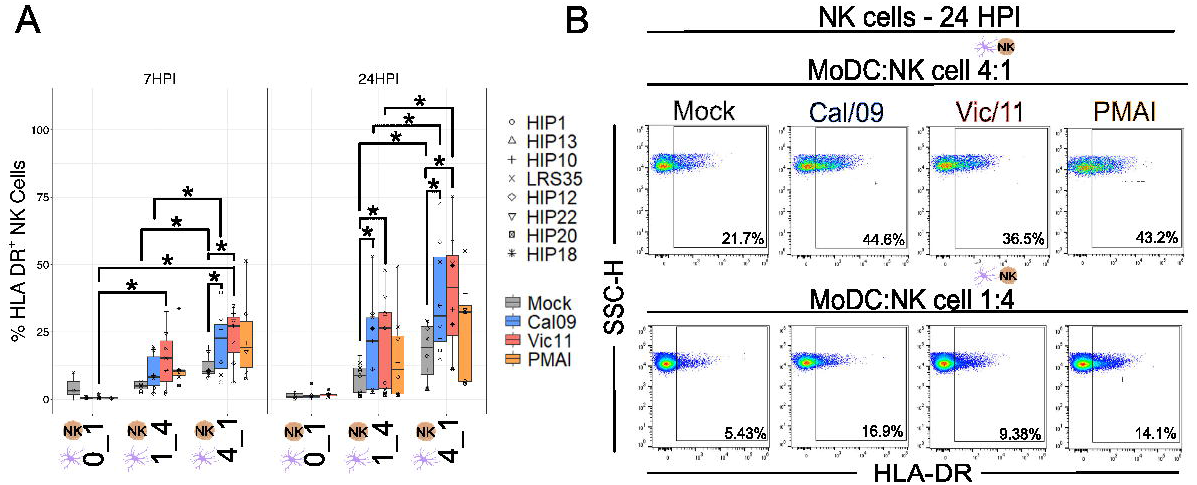
NK cells express HLA-DR after co-culture with IAV-infected MoDCs. **(A)** Summary plot of NK cell HLA-DR^+^ expression after culture alone or after 7 or 23 h co-culture with mock-treated or Cal/09- or Vic/11-infected MoDCs at an MOI of 3 (*n* = 8) as assessed by flow cytometry using an antibody specific to HLA-DR. In the NK cell-only condition (0:1), influenza virions were added at the same time as the NK cells. **(B)** Representative flow plot of HLA-DR expression on NK cells after 23 h co-culture with Cal/09- or Vic/11-infected MoDCs (MOI = 3) at a MoDC to NK cell ratio of 4:1 (top panel) or 1:4 (bottom panel). PMA/I treatment served as a positive control. \**p* < 0.05, Wilcoxon signed-rank test.

### Direct cell-cell contact is partially required for NK cell HLA-DR expression in co-culture with IAV-infected MoDCs

NK cells acquire HLA-DR either by transcriptional activation or by an intracellular membrane transfer process termed trogocytosis (42,43). Prior work has shown that MHC class II acquisition by NK cells from DCs through trogocytosis can be abrogated by preventing cell-to-cell contact using a 0.4 μm transwell insert with a semipermeable membrane (44). We therefore investigated whether cell-to-cell contact was necessary for NK cell HLA-DR expression in our culture system. To this end, NK cells were co-cultured with MoDCs either in direct contact or separated by a transwell. Elimination of direct cell contact led to a significant decrease in NK cell HLA-DR expression when cultured with either Cal/09- or Vic/11-infected MoDCs, however, not to baseline HLA-DR levels **(Figure 4A-B)**.

**Figure 4.**
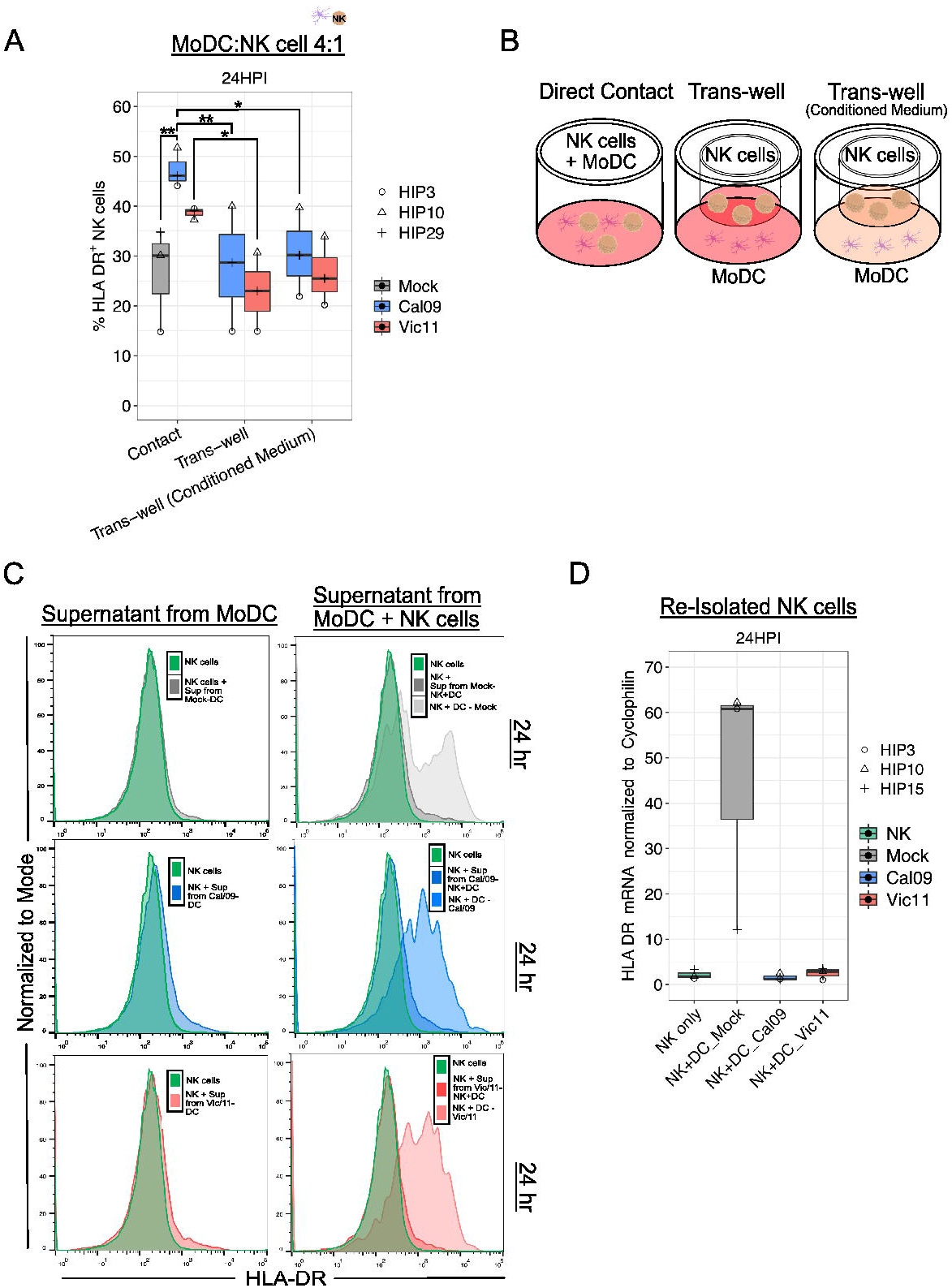
Direct contact is required for HLA-DR expression on NK cells after co-culture with IAV-infected MoDCs. **(A)** Percentage of HLA-DR^+^ NK cells after 23 h co-culture with mock-treated or IAV-infected MoDCs at an MOI of 3 as assessed by flow cytometry using an antibody specific to HLA-DR. DCs were cultured in direct contact with NK cells (contact), with DCs seeded in the dish and NK cells seeded in the trans-well with fresh medium (Trans-well) or with conditioned medium (Trans-well (conditioned medium) (*n* = 3). **(B)** Schematic of experimental set-up shown in **(A)**. **(C)** Representative flow histograms of the percentage of the maximum count of HLA-DR on NK cells after exposure to supernatant from mock-treated (top), Cal/09- (middle), or Vic/11- (bottom) infected MoDCs cultured alone (left panel) or with NK cells (right panel) at a MoDC to NK cell ratio of 4:1 (*n =* 3). **(D)** Relative HLA-DR mRNA levels in NK cells re-purified after 23 h co-culture with mock-treated or IAV-infected MoDCs when compared to NK cell control with RT-qPCR (*n =* 3). \**p* < 0.05, ns *p* > 0.05. Two-way ANOVA, Prism V9.0.0.

We considered the possibility that soluble mediators, such as cytokines synthesized due to receptor-ligand interactions, were also partly responsible for NK cell HLA-DR expression. To test this possibility, conditioned medium containing the soluble mediators harvested from previous IAV-infected MoDC-NK cell 24 h co-cultures were added to MoDCs and NK cells separated by the transwell. The addition of the conditioned medium failed to rescue NK cell HLA-DR expression to the level observed with direct cell-to-cell contact **(Figure 4A-B)**. Next, we determined whether cytokines produced by Cal/09- or Vic/11-infected MoDCs either alone or in co-culture with NK cells were sufficient to induce HLA-DR NK cell expression cultured in the absence of MoDCs. Transfer of supernatant from Cal/09- or Vic/11-infected MoDCs cultured in isolation **(Figure 4C; left panel)** or cultured with NK cells **(Figure 4C; right panel)** failed to induce the NK cell HLA-DR expression observed when cells were co-cultured (DC + NK condition; right panel). Finally, we next considered whether NK cells were also transcriptionally upregulating HLA-DR in addition to acquiring it from the DC membrane. To this end, NK cells co-cultured with mock or infected DCs were re-isolated and HLA-DR transcript levels were measured. NK cell interactions with IAV-infected DCs failed to induce HLA-DR transcripts, although induction was observed in NK cells exposed to mock-treated MoDCs **(Figure 4D)**. Collectively, these data are supportive of NK cells acquiring HLA-DR from the membrane of IAV-infected DCs via intercellular membrane transfer, independent of the cytokine milieu.

### MoDC-NK cell crosstalk increases the frequency of CD69^+^ T cells

We next used our co-culture system to investigate the role of MoDC-NK cell crosstalk on naïve T cell activation after MoDC IAV infection by using flow cytometry to evaluate the surface expression of CD69. CD69 is a classic early marker of lymphocyte activation, which functions to impair T cell egress from lymph nodes–likely to promote full T cell activation (45–47). To evaluate CD69 expression, MoDCs were either mock-treated or infected with Cal/09 or Vic/11 IAV strains followed by co-culture for 48 h with autologous NK cells and autologous naïve T cells. The 48 h time point was selected based on the finding that CD69 expression was at its peak between 18 and 48 h after anti-CD3 stimulation (48). PMA/I was used as a positive control for antigen-independent, chemical stimulation. Naïve T cells were cultured in four different conditions: independently, with MoDCs, with NK cells, or in triple co-culture at MoDC: NK cell: T cell ratios of 1:1:1 **(Figure 5A-B),** 4:1:1 **(Figure 5C-D)** and 1:4:1 **(Figure 5E-F)**. T cell expression of CD69 was highest under conditions that included both MoDCs and NK cells, with the highest percentages observed at the high MoDC to NK cell ratio (4:1) per T cell (4:1:1) **(Figure 5A-F)**. NK cells markedly increased T cell surface expression of CD69 when MoDCs were present under mock conditions or when MoDCs had been infected to either IAV strain **(Figure 5A-F)**. While CD69 median values decreased after MoDC exposure to IAV compared to mock treatment at 1:1:1 and 4:1:1 ratios, the decrease did not reach statistical significance. Collectively, these data support a model where early T cell activation by MoDCs is heightened by MoDC-NK cell crosstalk, under either steady state or IAV exposure conditions.

**Figure 5.**
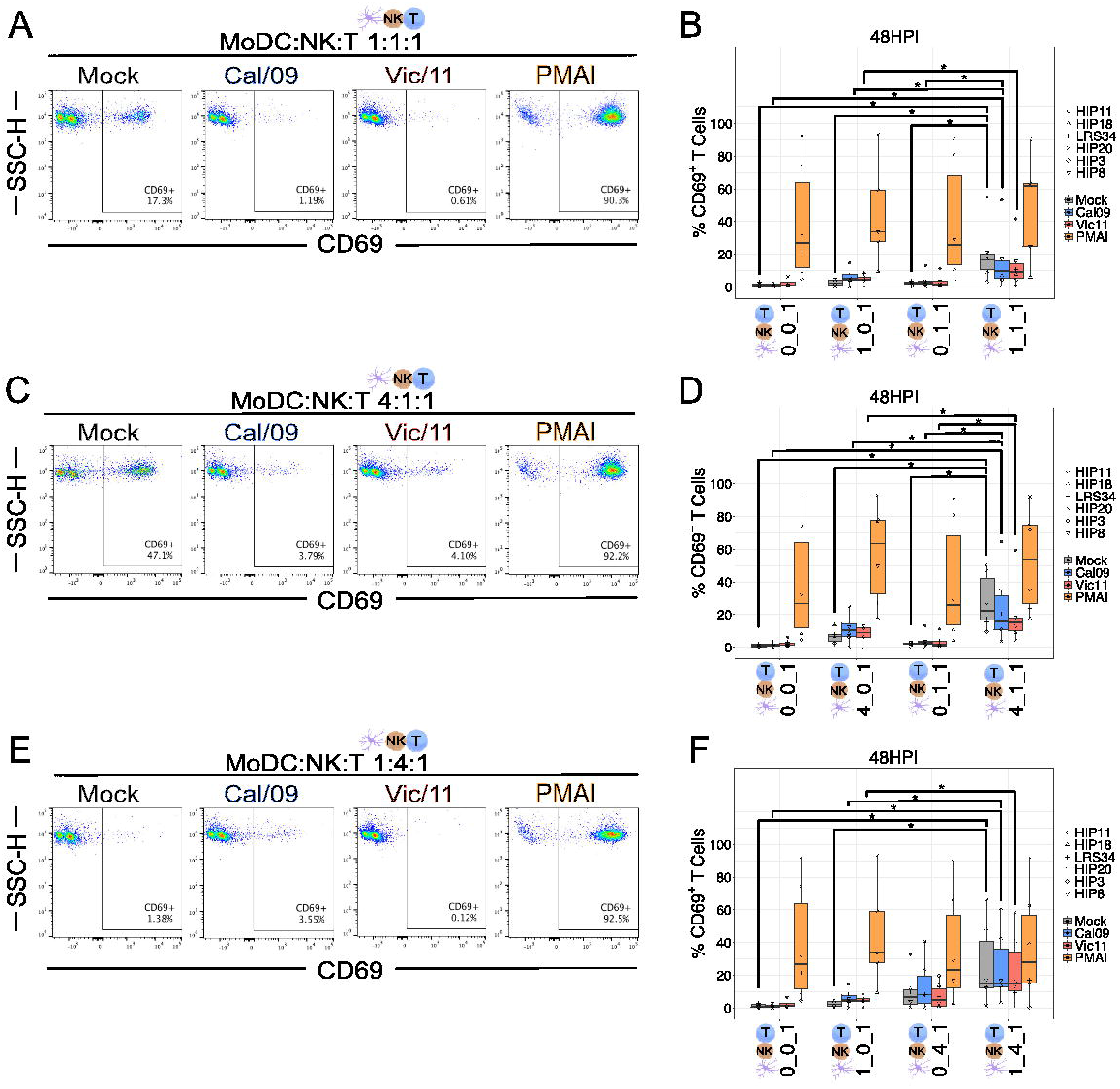
MoDC-NK cell crosstalk increases the frequency of an early marker of T cell activation. Representative flow plots of T cell CD69^+^ expression after 47 h co-culture with Cal/09- or Vic/11-infected MoDCs (MOI = 3) at a MoDC to NK cell to T cell ratio of **(A)** 1:1:1, **(C)** 4:1:1, and **(E)** 1:4:1. Summary plots of T cell CD69^+^ expression after culture with mock-treated or Cal/09 or Vic/11-infected MoDCs (MOI = 3) after 95 h co-culture with MoDCs and NK cells (*n* = 6) at a MoDC to NK cell to T cell ratio of **(B)** 1:1:1, **(D)** 4:1:1, and **(F)** 1:4:1 as assessed by flow cytometry using an antibody specific to CD25. PMA/Ionomycin (PMA/I) treatment served as a positive control. \**p* < 0.05, Wilcoxon signed-rank test.

### MoDC-NK cell crosstalk increases the frequency of CD25^+^ T cells

We then assessed the impact of MoDC-NK cell crosstalk and IAV infection by using flow cytometry to evaluate the surface expression of the alpha chain of the trimeric IL-2 receptor, CD25 (49). MoDCs were mock treated, infected with Cal/09 or Vic/11, or exposed to PMA/I. Naïve T cells were cultured in four different conditions: independently, with MoDCs, with NK cells, or in triple co-culture at MoDC: NK cell: T cell ratios of 1:1:1 **(Figure 6A-B),** 4:1:1 **(Figure 6C-D)** and 1:4:1 **(Figure 6E-F)**. The frequency of T cells expressing CD25 was highest under conditions that included both uninfected MoDCs and NK cells at 1:4 and 4:1 ratios, with the most modest increase observed at the MoDC to NK cell ratio of 1:1 per T cell **(Figure 6A-F)**. At a cell ratio of 1:1:1 **(Figure 6A-B)** and 1:4:1 **(Figure 6E-F)**, the percentage of CD25^+^ T cells significantly decreased after exposure to Cal/09 while at a cell ratio of 4:1:1, the percentage of CD25^+^ T cells significantly decreased after exposure to Vic/11 **(Figure 6C-D)**. No significant differences were measured between Cal/09 and Vic/11 strains. Increasing the ratio of MoDC to NK cell from 1:1 to 4:1 led to significantly higher T cell CD25 expression when MoDC were infected with Cal/09 (p = 0.04). Taken together, these data show that MoDC-NK cell crosstalk promotes higher levels of T cell CD25 expression, while IAV reduces the frequency of CD25^+^ T cells under the same conditions.

**Figure 6.**
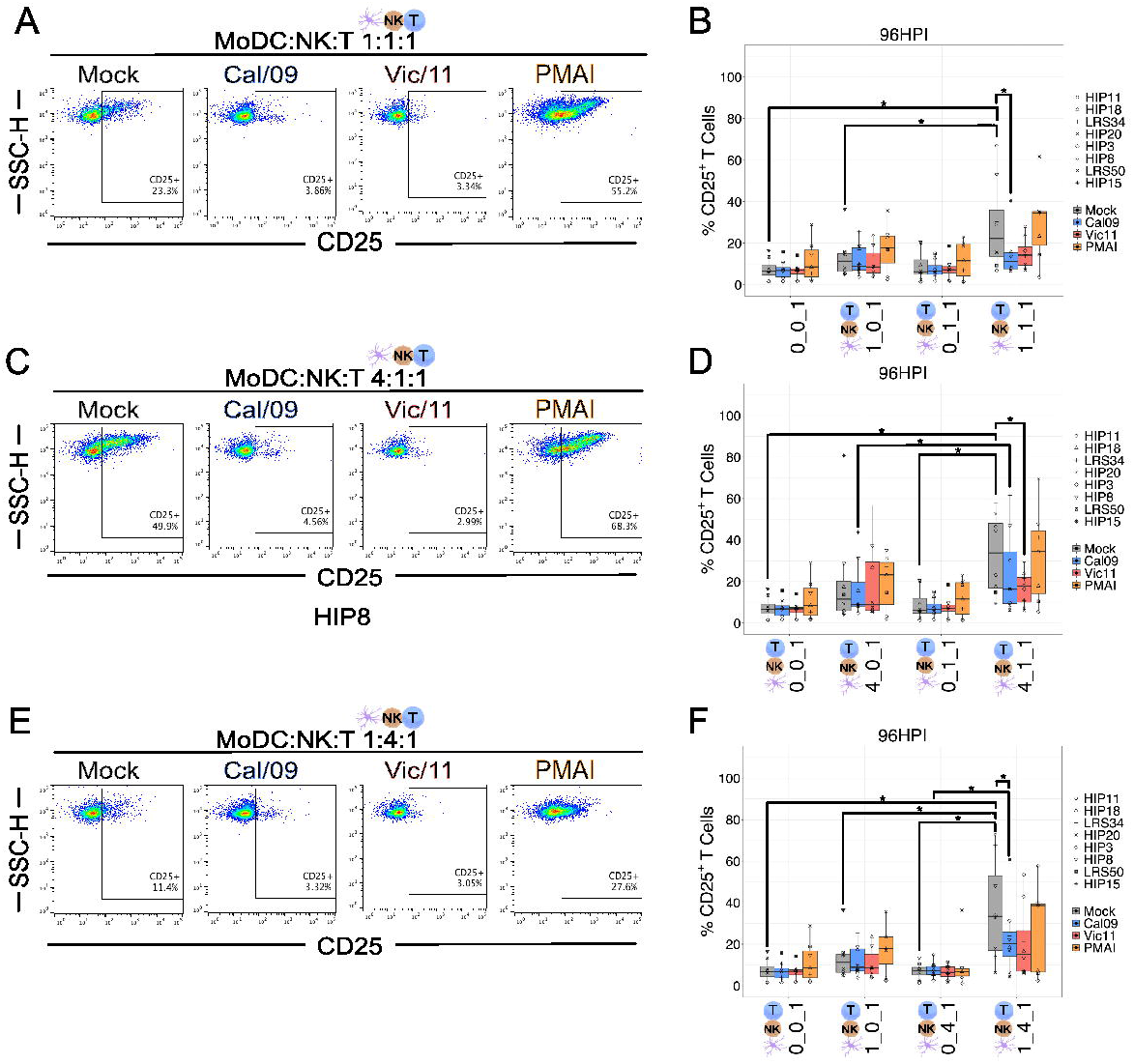
MoDC-NK cell crosstalk increases the frequency of a late marker of T cell activation. Representative flow plots of T cell CD25^+^ expression after 95 h co-culture with Cal/09- or Vic/11-infected MoDCs (MOI = 3) at a MoDC to NK cell to T cell ratio of **(A)** 1:1:1, **(C)** 4:1:1, and **(E)** 1:4:1. Summary plots of T cell CD25^+^ expression after culture with mock-treated or Cal/09 or Vic/11-infected MoDCs (MOI = 3) after 95 h co-culture with MoDCs and NK cells (*n* = 8) at a MoDC to NK cell to T cell ratio of **(B)** 1:1:1, **(D)** 4:1:1, and **(F)** 1:4:1 as assessed by flow cytometry using an antibody specific to CD25. PMA/Ionomycin (PMA/I) treatment served as a positive control. \**p* < 0.05, Wilcoxon signed-rank test.

### IAV infection leads to lower IFN-γ, Tumor Necrosis Factor (TNF), and IL-10 levels in MoDC-NK cell-T cell co-culture

The finding that surface markers of T cell activation were elevated when cultured with MoDCs and NK cells, yet were partially abrogated upon exposure to MoDC with IAV, led us to ask whether the cytokine milieu differed between mock and IAV-infected conditions depending on the cell types present. To this end, we used a custom MAGPIX panel to measure supernatant concentrations of the cytokines IFN-α2, IFN-γ, TNF, IL-2, IL-10, IL-12p70, IL-15, IL-18, and IL-21, **(Figure 7A-C, Figure S4,** and **Supplemental Table 1)**. These cytokines were selected based on their importance as key mediators of DC and NK cell crosstalk (50). T cells were either cultured in isolation (0:0:1), cultured with MoDCs (1:0:1), with NK cells (0:4:1) or with both MoDCs and NK cells (1:4:1). Influenza virions were included in 0:0:1 and 0:4:1 conditions. IFN-γ, TNF, and IL-10 were significantly increased in the 1:4:1 mock condition compared to the 0:0:1 (T cells alone) 1:0:1 (MoDCs and T cells), or 0:4:1 (NK cells and T cells) **(Figure 7A-C)**. Further, the mean fluorescence intensity of IFN-γ, TNF, and IL-10 was significantly decreased in the IAV-infected compared to uninfected samples in the 1:4:1 condition **(Figure 7A-C)**. Taken together, these results show that the interaction of MoDCs with NK cells profoundly modulates the cytokine milieu when cultured with T cells, leading to increased levels of IFN-γ, TNF, and IL-10, while MoDCs exposure to IAV curtails the production of these cytokines. The precise cellular source(s) of each cytokine in the co-culture system remains unknown and could be identified in future experiments using intracellular cytokine staining paired with lineage markers for DCs, NK cells, and T cells.

**Figure 7.**
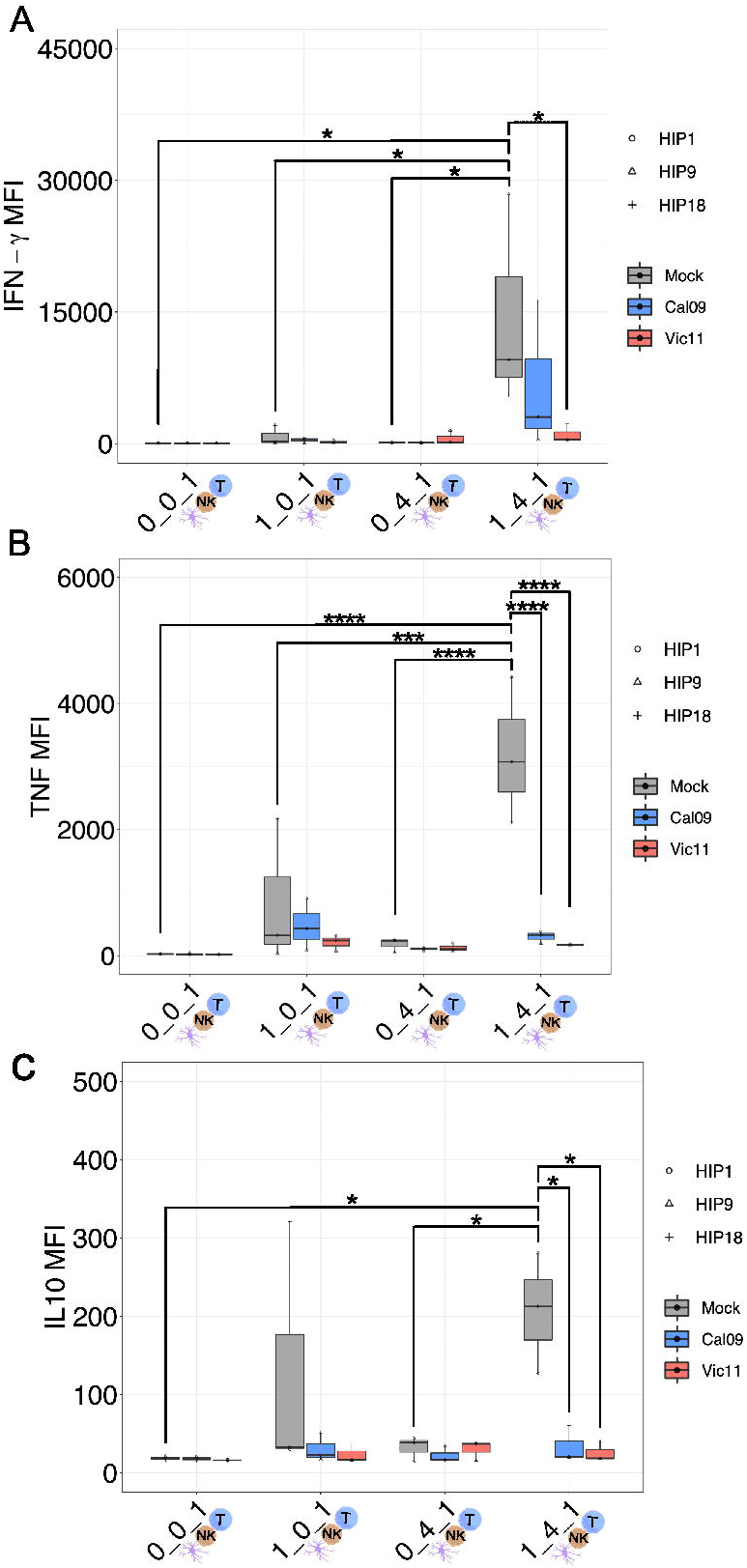
MoDC-NK cell crosstalk curtails the production of a broad array of cytokines as a result of influenza exposure. **(A)** MAGPIX data showing the mean fluorescence intensity (MFI) of **(A)** IFN-γ, **(B)** TNF, and **(C)** IL-10 present in the supernatant of either mock-treated or virion-exposed T cells (0:0:1), T cell co-cultured with mock or virus-infected MoDCs (MOI = 3, 96 HPI) (1:0:1), T cells co-cultured with mock or virion-exposed NK cells (0:4:1) or T cells co-cultured with both virus-infected and mock MoDCs (MOI = 3, 96 HPI) and NK cells (1:4:1) (*n* = 3). \**p* < 0.05, \*\**p* < 0.001, \*\*\**p* < 0.0001, \*\*\*\**p* < 0.0001. Two-way ANOVA, Prism V9.0.0.

## Discussion

In this study, we demonstrated that under steady-state conditions, DC-NK cell crosstalk contributes to T cell activation as documented by an increased expression of CD69 and CD25. Consistent with these findings, multi-plex cytokine analysis found elevated levels of IL-10, IFN-γ, and TNF under MoDC-NK cell-T cell culture conditions compared to T cells cultured alone or cultured with solely MoDCs or NK cells. While the cellular source of each cytokine remains to be defined, certain NK cell subsets express Galectin-1, which upon ligation of its receptor CD69 (seen to be upregulated on T cells by 48 h our co-cultures) triggers IL-10 secretion (51–53). Interestingly, while IL-10 is classically considered anti-inflammatory, this cytokine can also promote proliferation and cytotoxicity of both human CD8^+^ T cells and NK cells (54,55). NK cells are also a major source of IFN-γ and indeed, consistent with our results, a prior study by Ferlazzo et al. found that human NK cells secrete IFN-γ in response to both immature and mature DCs under steady-state conditions (56). Importantly, NK cells are also a significant source of IFN-γ following influenza virus vaccination in humans (57) and can regulate CD8^+^ T cell priming and DC migration in an IFN-γ-dependent manner in mice (58). NK cells and T cells are both able to release TNF and be the recipients of TNF activity, and this cytokine enhances the membrane expression of CD25 on activated T cells (59,60). While the precise role of NK cells in influencing the activation profile of CD8^+^ cytotoxic T cell and CD4^+^ helper cell subset has yet to be established, these findings provide insight into the role of DC-NK cell crosstalk in governing the downstream pan (CD3^+^) naïve T cell phenotype.

The second focus of this study was evaluating the role of MoDC-NK cell crosstalk in regulating pan T cell activation in response to a pandemic and a seasonal IAV strain. We observed that IAV significantly reduced the frequency of CD25^+^, although not CD69^+^, T cells when cultured with MoDCs and NK cells. This may represent a viral immune evasion strategy because a reduction in the number of high-affinity IL-2 receptors (CD25) could result in suboptimal pathogen-specific T cell expansion and memory responses (61). Indeed, other studies have demonstrated that IL-2 signaling is essential for the development of robust secondary memory CD8^+^ T cell responses during viral infection (62). Compatible with the lower levels of T cell surface markers of activation, IAV exposure reduced MoDC expression of CD83 and CD86 molecules and curtailed cytokine release of IL-10, IFN-γ, and TNF in the triple co-culture condition. NK cell-mediated DC maturation is dependent on TNF, therefore lower levels observed after IAV exposure could potentially hinder DC activation and subsequent stimulation of naïve T cells (33). The two IAV strains used in this study; A/California/07/2009 (Cal/09: H1N1) and A/Victoria/361/2011 (Vic/11; H3N2), exhibit qualitatively distinct responses on NK cell IFN-γ cytokine production, and indeed elevated levels of the T_H_1 signature cytokine IFN-γ were detected in the Cal/09 conditions compared to Vic/11 triple co-culture conditions (**Figure 7A**) (31). While both IAV strains (Cal/09 or Vic/11) impact the expression of T cell surface activation markers to a similar extent, future studies are needed to investigate whether Cal/09 polarizes T cells responses towards T_H_1 CD4^+^ T cells – critical for the generation of the cytotoxic T cells needed to eliminate virus-infected cells and promote disease resolution–compared to Vic/11 (5,63).

NK cells can switch between functional states including exerting their hallmark anti-viral and anti-tumor cytotoxic effector functions and providing a regulatory role in dampening inflammatory immune responses (64). This functional plasticity is conferred, in part, by the *de novo* expression or upregulation of activation markers such as CD69, CD25, NKp44, CD16, and HLA-DR (65–68). Indeed, in our study, we found that IAV exposure of MoDC followed by co-culture with NK cells led to an expansion in HLA-DR^+^ NK cells. We thus explored the mechanism responsible for the emergence of the HLA-DR^+^ NK cell subset after exposure to IAV-exposed MoDCs. We found that NK cell HLA-DR expression was reduced if direct cell-to-cell contact was prevented, consistent with prior work that found that NK cells can acquire HLA-DR from DCs through intercellular membrane transfer called “trogocytosis” (44). Prior reports have shown that cytokines, including IL-2, IL-15, IL-18, and IL-21 can promote differentiation of an HLA-DR-expressing NK cell subset (57), although the addition of supernatant from IAV-infected MoDCs alone or in co-culture with NK cells failed to stimulate HLA-DR expression in our culture system (69). This is consistent with our multi-plex cytokine findings, which did not show significantly increased levels of these cytokines in IAV-infected conditions compared to mock treatment **(Figure S4** and **Table S1)**. Further, while HLA-DR transcripts were induced in NK cells exposed to uninfected MoDCs, they were not induced by IAV-infected MoDCs, suggesting that the HLA-DR^+^ NK cell subset expanded by acquiring HLA-DR from the surface of IAV-infected MoDCs.

Interestingly, prior work in a murine model found that HLA-DR^+^ NK cells can inhibit DC-induced CD4^+^ T cell responses by competitive antigen presentation, possibly due to insufficient expression of costimulatory molecules (44). Similarly, we found that NK cells co-cultured with IAV-infected MoDCs failed to significantly upregulate either CD83 or CD86 costimulatory molecules **(Figure S3)**. This raises the question of whether this human HLA-DR^+^ NK cell subset is involved in reducing IAV-mediated human T cell activation; efforts to investigate the functional role of this subset are underway. HLA-DR^+^ NK cell subsets have been found in both humans and mice and present under steady-state conditions and a range of diseases (40). For example, an HLA-DR^+^ NK cell subset was found to be expanded in the peripheral blood of patients with primary tuberculosis (41,70). The CD56^dim^ NK cell subset from HIV-infected individuals displayed elevated HLA-DR expression and impaired IFN-γ production compared to uninfected controls (71). Here we provide the first report that IAV-infected MoDCs cultured with NK cells lead to an increase in human NK cell HLA-DR expression in the context of influenza infection *ex vivo*.

One potential limitation of this study is that the results were generated under *ex vivo* experimental conditions. *In vivo*, influenza viral infection may impact DC-NK cell interactions. For example, IAV infection triggers blood DC subsets to secrete chemokines (CXCL16, CXCL1, CXCL2, and CXCL3) for which NK cells have the cognate receptors (CXCR6, CXCR2) and are thus capable of potentially attracting NK cells to participate in crosstalk (72–74). Further, virally infected plasmacytoid DCs produce CCL4 and CXCL10 that are capable of recruiting NK cells in chemotaxis assays (74). Isolation of PBMCs directly from healthy donors followed by isolation of autologous NK cells and differentiation of MoDCs from monocytes allows for investigation of the impact of DC and NK cell interactions in an *ex vivo* setting. Analysis of immune cells harvested from patients acutely infected with influenza, for example from bronchial lavage fluid, peripheral blood, or lymph nodes, would be of interest to evaluate whether the observed expression patterns are seen *in vivo*. A second limitation is that our system investigated only one DC subtype, which fails to recapitulate the diversity of DC subsets found *in vivo* (75). MoDCs were selected because circulating monocytes serve as a major precursor for antigen-presenting DCs within peripheral tissues including the lung (76–78). MoDCs formed *de novo* at sites of infection efficiently capture antigen, migrate to local lymph nodes, and effectively prime and cross-prime T cells to generate pathogen-specific immunity (79–81). Further, they have become an attractive target for vaccine design because unlike blood DCs which comprise a very small fraction of circulating blood cells (<1%), large numbers of monocytes can be easily obtained from whole blood samples (82). Beyond the ease of collection and experimental manipulation, *in vivo* studies have also demonstrated that MoDCs are important during microbial infection and specifically in cross-priming T cells to generate pathogen-specific immunity (79,81). Future studies to investigate the role of diverse DC subtypes are warranted and, along these lines, an allogenic plasmacytoid DC cell line has shown promising clinical observations in patients with metastatic stage IV melanoma (83).

In summary, our findings demonstrate first that NK cells can significantly impact DC-mediated T cell activation and second that IAV induces a tolerogenic-like activation state in MoDCs, contributing to abrogated T cell activation. Understanding the immune pathways that govern DC-mediated T cell responses may have future therapeutic implications. Along these lines, in the United States, the FDA has approved Sipuleucel-T, a DC-based vaccine that confers a survival benefit for patients with metastatic prostate cancer (84,85). However, a major barrier to this type of therapeutic option includes regulation of the DC functional state. Overall, our human *ex vivo* data may help inform the signals required to elicit an optimal DC functional state, improving the immunogenicity and efficacy of DC-based immunotherapies and vaccines. Determining whether such approaches could be leveraged to manipulate T cells to stimulate protective immunity or limit immunopathology during infection are key future endeavors.

## Author Contributions

Conceptualization: L.M.K and B.L.; Experiments: C.U, A.G., A.H. J.G. M.F. S.M. S.R. Analyses: L.M.K; Writing: L.M.K., B.L., Funding L.M.K, and B.L.

## Nomenclature

Dendritic cells (DC); Natural Killer cells (NK cells); monocyte-derived dendritic cell (MoDC); Influenza A Virus (IAV), and Peripheral blood mononuclear cells (PBMCs).

## Acknowledgments

We would like to thank Emma Parent and Connor Morson for their careful reading of the manuscript, Dr. Jeremy Ellermeier for helpful discussions, and the study participants.

