## Supplemental Material for "Dendritic cell-Natural Killer cell Crosstalk Modulates T cell activation in Response to Influenza A Viral Infection"

Supplementary Material

### Supplementary Figures

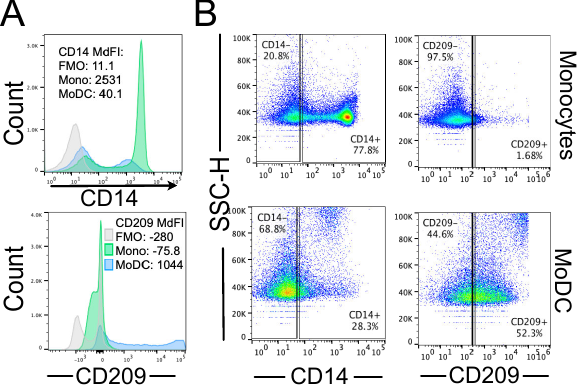

**Supplementary Figure 1. Differentiation of monocytes into MoDCs. (A)** Representative histograms comparing fluorescence minus one control (FMO) of CD14 (top) expression and CD209 expression (bottom) on monocytes before and after IL-4 and GM-CSF differentiation into MoDCs. **(B)** Representative flow plots of CD14 and CD209 expression on monocytes (top) before and after IL-4 and GM-CSF differentiation into MoDCs (bottom).

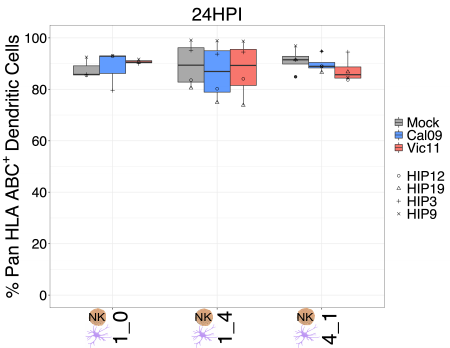

**Supplementary Figure 2. Expression of HLA class I molecules on MoDCs after IAV exposure.** HLA Class I A, B, and C expression on MoDCs either alone (0:1) or after co-culture with NK cells at 1:4 or 4:1 (MoDC: NK cell) ratios for 23 h with either Cal/09- or Vic/11-infected MoDCs at an MOI of 3 (*n* = 4).

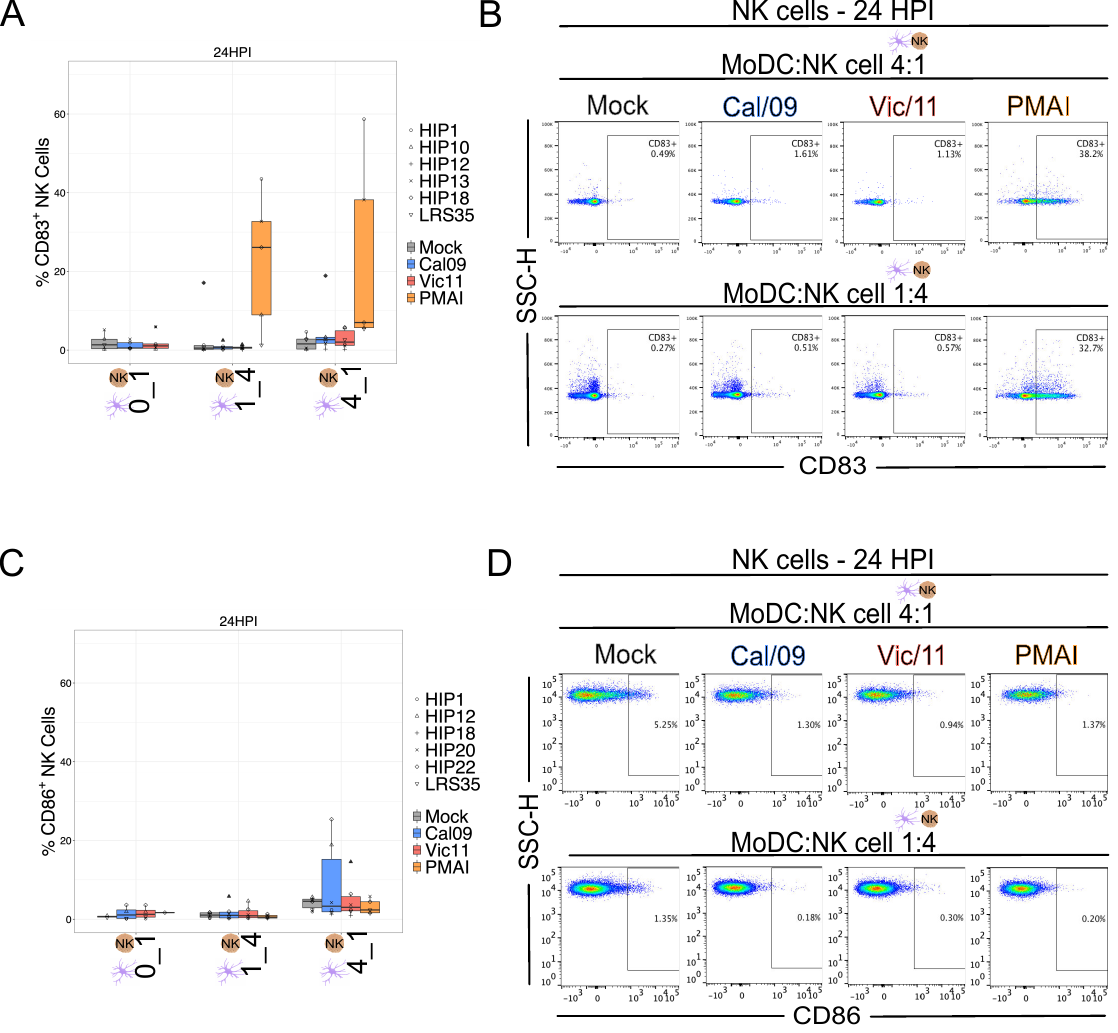

**Supplementary Figure 3. NK cells do not significantly upregulate CD83 or CD86 after co-culture with IAV-infected MoDCs. (A)** Summary plot of NK cell CD83^+^ expression after 23 h co-culture with mock-treated or Cal/09 or Vic/11-infected MoDCs (MOI = 3) (*n* = 6) as assessed by flow cytometry using an antibody specific to CD83. **(B)** Representative flow plot of CD83 expression on NK cells after 23 h co-culture with Cal/09- or Vic/11-infected MoDCs (MOI = 3) at a MoDC to NK cell ratio of 4:1 (top panel) or 1:4 (bottom panel). **(C)** Summary plot of NK cell CD86^+^ expression after 23 h co-culture with mock-treated or Cal/09- or Vic/11-infected MoDCs (MOI = 3) (*n* = 6) as assessed by flow cytometry using an antibody specific to CD86. (**D**) Representative flow plot of CD86 expression on NK cells after 23 h co-culture with Cal/09- or Vic/11-infected MoDCs (MOI = 3) at a MoDC to NK cell ratio of 4:1 (top panel) or 1:4 (bottom panel). PMA/I treatment served as a positive control.

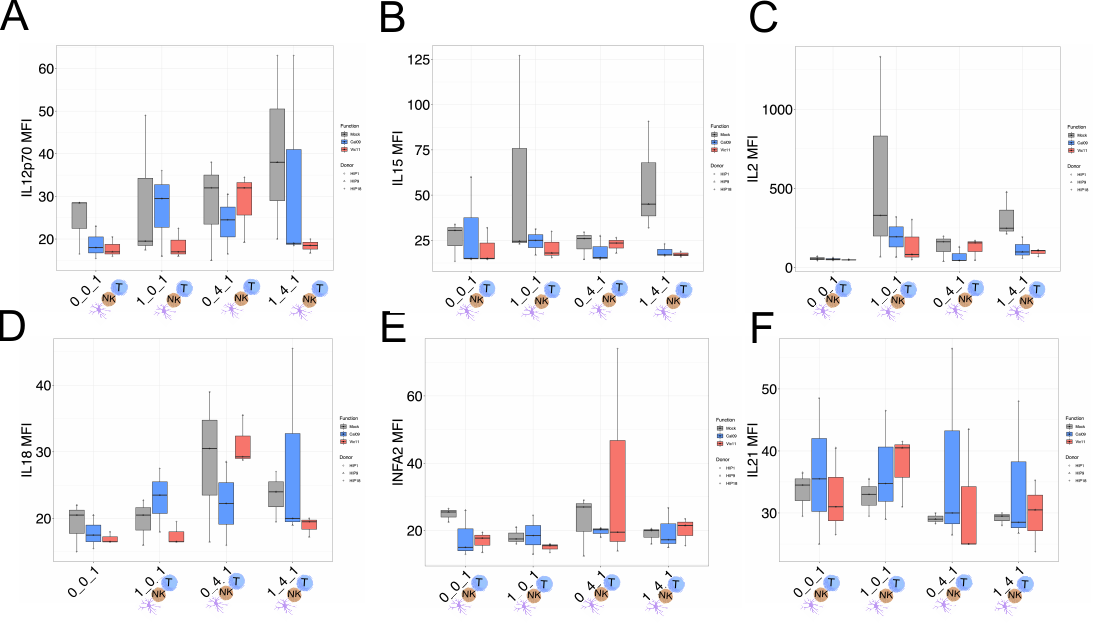

**Supplementary Figure 4. Quantification of six cytokines in supernatants harvested 96 h post-IAV infection from MoDC-NK cell-T cell co-cultures.** MAGPIX data showing the mean fluorescence intensity (MFI) of cytokines **(A)** IL-12p70, **(B)** IL-15, **(C)** IL-2, **(D)** IL-18, (**E)** IFN-α2, **(F)** and IL-21 present in the supernatant of either mock-treated or virion-exposed T cells (0:0:1), T cell co-cultured with mock or virus-infected MoDCs (MOI = 3, 96 HPI) (1:0:1), T cells co-cultured with mock or virion-exposed NK cells (0:4:1) or T cells co-cultured with mock or virus-infected MoDCs (MOI = 3, 96 HPI) and NK cells (1:4:1).

**Supplementary Table 1. Raw MFI values of cytokines in supernatants harvested 96 h post-IAV infection from MoDC-NK cell-T cell co-cultures quantified by MAGPIX.**

| Donor | Treatment | Condition | Cytokine | Value (MFI) |
| --- | --- | --- | --- | --- |
| HIP1 | Mock | 1_4_1 | IFNA2 | 16 |
| HIP1 | Mock | 1_0_1 | IFNA2 | 17.5 |
| HIP1 | Mock | 0_4_1 | IFNA2 | 29 |
| HIP1 | Mock | 0_0_1 | IFNA2 | 22.5 |
| HIP1 | Cal09 | 1_4_1 | IFNA2 | 26.75 |
| HIP1 | Cal09 | 1_0_1 | IFNA2 | 24.5 |
| HIP1 | Cal09 | 0_4_1 | IFNA2 | 18 |
| HIP1 | Cal09 | 0_0_1 | IFNA2 | 15 |
| HIP1 | Vic11 | 1_4_1 | IFNA2 | 23.5 |
| HIP1 | Vic11 | 1_0_1 | IFNA2 | 15.5 |
| HIP1 | Vic11 | 0_4_1 | IFNA2 | 19.5 |
| HIP1 | Vic11 | 0_0_1 | IFNA2 | 17.75 |
| HIP9 | Mock | 1_4_1 | IFNA2 | 20.5 |
| HIP9 | Mock | 1_0_1 | IFNA2 | 21 |
| HIP9 | Mock | 0_4_1 | IFNA2 | 12.5 |
| HIP9 | Mock | 0_0_1 | IFNA2 | 25.5 |
| HIP9 | Cal09 | 1_4_1 | IFNA2 | 15 |
| HIP9 | Cal09 | 1_0_1 | IFNA2 | 13 |
| HIP9 | Cal09 | 0_4_1 | IFNA2 | 20.75 |
| HIP9 | Cal09 | 0_0_1 | IFNA2 | 26 |
| HIP9 | Vic11 | 1_4_1 | IFNA2 | 21.5 |
| HIP9 | Vic11 | 1_0_1 | IFNA2 | 16 |
| HIP9 | Vic11 | 0_4_1 | IFNA2 | 14 |
| HIP9 | Vic11 | 0_0_1 | IFNA2 | 13.5 |
| HIP18 | Mock | 1_4_1 | IFNA2 | 20 |
| HIP18 | Mock | 1_0_1 | IFNA2 | 16 |
| HIP18 | Mock | 0_4_1 | IFNA2 | 27 |
| HIP18 | Mock | 0_0_1 | IFNA2 | 26.5 |
| HIP18 | Cal09 | 1_4_1 | IFNA2 | 17.25 |
| HIP18 | Cal09 | 1_0_1 | IFNA2 | 18.5 |
| HIP18 | Cal09 | 0_4_1 | IFNA2 | 20.25 |
| HIP18 | Cal09 | 0_0_1 | IFNA2 | 13 |
| HIP18 | Vic11 | 1_4_1 | IFNA2 | 15.5 |
| HIP18 | Vic11 | 1_0_1 | IFNA2 | 13.5 |
| HIP18 | Vic11 | 0_4_1 | IFNA2 | 74 |
| HIP18 | Vic11 | 0_0_1 | IFNA2 | 19.5 |
| HIP1 | Mock | 1_4_1 | IFNG | 9560 |
| HIP1 | Mock | 1_0_1 | IFNG | 229.5 |
| HIP1 | Mock | 0_4_1 | IFNG | 143.5 |
| HIP1 | Mock | 0_0_1 | IFNG | 47 |
| HIP1 | Cal09 | 1_4_1 | IFNG | 3030 |
| HIP1 | Cal09 | 1_0_1 | IFNG | 518 |
| HIP1 | Cal09 | 0_4_1 | IFNG | 65.5 |
| HIP1 | Cal09 | 0_0_1 | IFNG | 48 |
| HIP1 | Vic11 | 1_4_1 | IFNG | 444.5 |
| HIP1 | Vic11 | 1_0_1 | IFNG | 381 |
| HIP1 | Vic11 | 0_4_1 | IFNG | 149 |
| HIP1 | Vic11 | 0_0_1 | IFNG | 45 |
| HIP9 | Mock | 1_4_1 | IFNG | 28414 |
| HIP9 | Mock | 1_0_1 | IFNG | 44 |
| HIP9 | Mock | 0_4_1 | IFNG | 43 |
| HIP9 | Mock | 0_0_1 | IFNG | 58.5 |
| HIP9 | Cal09 | 1_4_1 | IFNG | 450 |
| HIP9 | Cal09 | 1_0_1 | IFNG | 52 |
| HIP9 | Cal09 | 0_4_1 | IFNG | 89.5 |
| HIP9 | Cal09 | 0_0_1 | IFNG | 42 |
| HIP9 | Vic11 | 1_4_1 | IFNG | 438.5 |
| HIP9 | Vic11 | 1_0_1 | IFNG | 106 |
| HIP9 | Vic11 | 0_4_1 | IFNG | 1516 |
| HIP9 | Vic11 | 0_0_1 | IFNG | 47 |
| HIP18 | Mock | 1_4_1 | IFNG | 5516 |
| HIP18 | Mock | 1_0_1 | IFNG | 2078 |
| HIP18 | Mock | 0_4_1 | IFNG | 120 |
| HIP18 | Mock | 0_0_1 | IFNG | 41 |
| HIP18 | Cal09 | 1_4_1 | IFNG | 16225 |
| HIP18 | Cal09 | 1_0_1 | IFNG | 621 |
| HIP18 | Cal09 | 0_4_1 | IFNG | 100 |
| HIP18 | Cal09 | 0_0_1 | IFNG | 43 |
| HIP18 | Vic11 | 1_4_1 | IFNG | 2247 |
| HIP18 | Vic11 | 1_0_1 | IFNG | 81 |
| HIP18 | Vic11 | 0_4_1 | IFNG | 106 |
| HIP18 | Vic11 | 0_0_1 | IFNG | 47 |
| HIP1 | Mock | 1_4_1 | IL2 | 212.5 |
| HIP1 | Mock | 1_0_1 | IL2 | 1333 |
| HIP1 | Mock | 0_4_1 | IL2 | 197.5 |
| HIP1 | Mock | 0_0_1 | IL2 | 55.5 |
| HIP1 | Cal09 | 1_4_1 | IL2 | 193.5 |
| HIP1 | Cal09 | 1_0_1 | IL2 | 320.5 |
| HIP1 | Cal09 | 0_4_1 | IL2 | 44.75 |
| HIP1 | Cal09 | 0_0_1 | IL2 | 51.5 |
| HIP1 | Vic11 | 1_4_1 | IL2 | 107 |
| HIP1 | Vic11 | 1_0_1 | IL2 | 303.5 |
| HIP1 | Vic11 | 0_4_1 | IL2 | 170.5 |
| HIP1 | Vic11 | 0_0_1 | IL2 | 47.75 |
| HIP9 | Mock | 1_4_1 | IL2 | 477.75 |
| HIP9 | Mock | 1_0_1 | IL2 | 69.25 |
| HIP9 | Mock | 0_4_1 | IL2 | 39.25 |
| HIP9 | Mock | 0_0_1 | IL2 | 70.5 |
| HIP9 | Cal09 | 1_4_1 | IL2 | 98.75 |
| HIP9 | Cal09 | 1_0_1 | IL2 | 66.75 |
| HIP9 | Cal09 | 0_4_1 | IL2 | 131 |
| HIP9 | Cal09 | 0_0_1 | IL2 | 62.5 |
| HIP9 | Vic11 | 1_4_1 | IL2 | 111.5 |
| HIP9 | Vic11 | 1_0_1 | IL2 | 83 |
| HIP9 | Vic11 | 0_4_1 | IL2 | 46.5 |
| HIP9 | Vic11 | 0_0_1 | IL2 | 51.75 |
| HIP18 | Mock | 1_4_1 | IL2 | 248.5 |
| HIP18 | Mock | 1_0_1 | IL2 | 330 |
| HIP18 | Mock | 0_4_1 | IL2 | 163 |
| HIP18 | Mock | 0_0_1 | IL2 | 43.5 |
| HIP18 | Cal09 | 1_4_1 | IL2 | 60 |
| HIP18 | Cal09 | 1_0_1 | IL2 | 194.5 |
| HIP18 | Cal09 | 0_4_1 | IL2 | 44 |
| HIP18 | Cal09 | 0_0_1 | IL2 | 45.25 |
| HIP18 | Vic11 | 1_4_1 | IL2 | 70 |
| HIP18 | Vic11 | 1_0_1 | IL2 | 49.5 |
| HIP18 | Vic11 | 0_4_1 | IL2 | 156 |
| HIP18 | Vic11 | 0_0_1 | IL2 | 48.5 |
| HIP1 | Mock | 1_4_1 | IL10 | 126.75 |
| HIP1 | Mock | 1_0_1 | IL10 | 30 |
| HIP1 | Mock | 0_4_1 | IL10 | 44 |
| HIP1 | Mock | 0_0_1 | IL10 | 19 |
| HIP1 | Cal09 | 1_4_1 | IL10 | 20.5 |
| HIP1 | Cal09 | 1_0_1 | IL10 | 22.5 |
| HIP1 | Cal09 | 0_4_1 | IL10 | 16.5 |
| HIP1 | Cal09 | 0_0_1 | IL10 | 18 |
| HIP1 | Vic11 | 1_4_1 | IL10 | 18.5 |
| HIP1 | Vic11 | 1_0_1 | IL10 | 16.5 |
| HIP1 | Vic11 | 0_4_1 | IL10 | 38 |
| HIP1 | Vic11 | 0_0_1 | IL10 | 17 |
| HIP9 | Mock | 1_4_1 | IL10 | 280.5 |
| HIP9 | Mock | 1_0_1 | IL10 | 32.5 |
| HIP9 | Mock | 0_4_1 | IL10 | 14 |
| HIP9 | Mock | 0_0_1 | IL10 | 21 |
| HIP9 | Cal09 | 1_4_1 | IL10 | 19.25 |
| HIP9 | Cal09 | 1_0_1 | IL10 | 16.5 |
| HIP9 | Cal09 | 0_4_1 | IL10 | 34 |
| HIP9 | Cal09 | 0_0_1 | IL10 | 20.5 |
| HIP9 | Vic11 | 1_4_1 | IL10 | 17.75 |
| HIP9 | Vic11 | 1_0_1 | IL10 | 15.5 |
| HIP9 | Vic11 | 0_4_1 | IL10 | 15.5 |
| HIP9 | Vic11 | 0_0_1 | IL10 | 16 |
| HIP18 | Mock | 1_4_1 | IL10 | 212.5 |
| HIP18 | Mock | 1_0_1 | IL10 | 321 |
| HIP18 | Mock | 0_4_1 | IL10 | 38.5 |
| HIP18 | Mock | 0_0_1 | IL10 | 15 |
| HIP18 | Cal09 | 1_4_1 | IL10 | 60.5 |
| HIP18 | Cal09 | 1_0_1 | IL10 | 51 |
| HIP18 | Cal09 | 0_4_1 | IL10 | 16 |
| HIP18 | Cal09 | 0_0_1 | IL10 | 15 |
| HIP18 | Vic11 | 1_4_1 | IL10 | 41 |
| HIP18 | Vic11 | 1_0_1 | IL10 | 39 |
| HIP18 | Vic11 | 0_4_1 | IL10 | 37 |
| HIP18 | Vic11 | 0_0_1 | IL10 | 16 |
| HIP1 | Mock | 1_4_1 | IL12p70 | 20 |
| HIP1 | Mock | 1_0_1 | IL12p70 | 19.5 |
| HIP1 | Mock | 0_4_1 | IL12p70 | 38 |
| HIP1 | Mock | 0_0_1 | IL12p70 | 28.5 |
| HIP1 | Cal09 | 1_4_1 | IL12p70 | 18.5 |
| HIP1 | Cal09 | 1_0_1 | IL12p70 | 29.5 |
| HIP1 | Cal09 | 0_4_1 | IL12p70 | 16.5 |
| HIP1 | Cal09 | 0_0_1 | IL12p70 | 18 |
| HIP1 | Vic11 | 1_4_1 | IL12p70 | 18.5 |
| HIP1 | Vic11 | 1_0_1 | IL12p70 | 17 |
| HIP1 | Vic11 | 0_4_1 | IL12p70 | 32 |
| HIP1 | Vic11 | 0_0_1 | IL12p70 | 20.5 |
| HIP9 | Mock | 1_4_1 | IL12p70 | 38 |
| HIP9 | Mock | 1_0_1 | IL12p70 | 17.5 |
| HIP9 | Mock | 0_4_1 | IL12p70 | 15 |
| HIP9 | Mock | 0_0_1 | IL12p70 | 28.5 |
| HIP9 | Cal09 | 1_4_1 | IL12p70 | 19 |
| HIP9 | Cal09 | 1_0_1 | IL12p70 | 16 |
| HIP9 | Cal09 | 0_4_1 | IL12p70 | 30.5 |
| HIP9 | Cal09 | 0_0_1 | IL12p70 | 23 |
| HIP9 | Vic11 | 1_4_1 | IL12p70 | 16.75 |
| HIP9 | Vic11 | 1_0_1 | IL12p70 | 16 |
| HIP9 | Vic11 | 0_4_1 | IL12p70 | 19.25 |
| HIP9 | Vic11 | 0_0_1 | IL12p70 | 17 |
| HIP18 | Mock | 1_4_1 | IL12p70 | 63 |
| HIP18 | Mock | 1_0_1 | IL12p70 | 49 |
| HIP18 | Mock | 0_4_1 | IL12p70 | 32 |
| HIP18 | Mock | 0_0_1 | IL12p70 | 16.5 |
| HIP18 | Cal09 | 1_4_1 | IL12p70 | 63 |
| HIP18 | Cal09 | 1_0_1 | IL12p70 | 36 |
| HIP18 | Cal09 | 0_4_1 | IL12p70 | 24.5 |
| HIP18 | Cal09 | 0_0_1 | IL12p70 | 15.5 |
| HIP18 | Vic11 | 1_4_1 | IL12p70 | 20 |
| HIP18 | Vic11 | 1_0_1 | IL12p70 | 22.5 |
| HIP18 | Vic11 | 0_4_1 | IL12p70 | 34.5 |
| HIP18 | Vic11 | 0_0_1 | IL12p70 | 16 |
| HIP1 | Mock | 1_4_1 | IL15 | 32 |
| HIP1 | Mock | 1_0_1 | IL15 | 23 |
| HIP1 | Mock | 0_4_1 | IL15 | 29.5 |
| HIP1 | Mock | 0_0_1 | IL15 | 30.5 |
| HIP1 | Cal09 | 1_4_1 | IL15 | 16.5 |
| HIP1 | Cal09 | 1_0_1 | IL15 | 31.25 |
| HIP1 | Cal09 | 0_4_1 | IL15 | 14.5 |
| HIP1 | Cal09 | 0_0_1 | IL15 | 15 |
| HIP1 | Vic11 | 1_4_1 | IL15 | 17 |
| HIP1 | Vic11 | 1_0_1 | IL15 | 15.5 |
| HIP1 | Vic11 | 0_4_1 | IL15 | 26.5 |
| HIP1 | Vic11 | 0_0_1 | IL15 | 32 |
| HIP9 | Mock | 1_4_1 | IL15 | 45 |
| HIP9 | Mock | 1_0_1 | IL15 | 24.5 |
| HIP9 | Mock | 0_4_1 | IL15 | 14.5 |
| HIP9 | Mock | 0_0_1 | IL15 | 33.75 |
| HIP9 | Cal09 | 1_4_1 | IL15 | 17 |
| HIP9 | Cal09 | 1_0_1 | IL15 | 17 |
| HIP9 | Cal09 | 0_4_1 | IL15 | 27.5 |
| HIP9 | Cal09 | 0_0_1 | IL15 | 60 |
| HIP9 | Vic11 | 1_4_1 | IL15 | 16 |
| HIP9 | Vic11 | 1_0_1 | IL15 | 18 |
| HIP9 | Vic11 | 0_4_1 | IL15 | 18 |
| HIP9 | Vic11 | 0_0_1 | IL15 | 15 |
| HIP18 | Mock | 1_4_1 | IL15 | 90.75 |
| HIP18 | Mock | 1_0_1 | IL15 | 127 |
| HIP18 | Mock | 0_4_1 | IL15 | 26 |
| HIP18 | Mock | 0_0_1 | IL15 | 13.5 |
| HIP18 | Cal09 | 1_4_1 | IL15 | 23 |
| HIP18 | Cal09 | 1_0_1 | IL15 | 25 |
| HIP18 | Cal09 | 0_4_1 | IL15 | 15.5 |
| HIP18 | Cal09 | 0_0_1 | IL15 | 14.5 |
| HIP18 | Vic11 | 1_4_1 | IL15 | 19 |
| HIP18 | Vic11 | 1_0_1 | IL15 | 30 |
| HIP18 | Vic11 | 0_4_1 | IL15 | 23.5 |
| HIP18 | Vic11 | 0_0_1 | IL15 | 14.5 |
| HIP1 | Mock | 1_4_1 | IL18 | 19.5 |
| HIP1 | Mock | 1_0_1 | IL18 | 20.5 |
| HIP1 | Mock | 0_4_1 | IL18 | 39 |
| HIP1 | Mock | 0_0_1 | IL18 | 20.5 |
| HIP1 | Cal09 | 1_4_1 | IL18 | 20 |
| HIP1 | Cal09 | 1_0_1 | IL18 | 23.5 |
| HIP1 | Cal09 | 0_4_1 | IL18 | 22.25 |
| HIP1 | Cal09 | 0_0_1 | IL18 | 17.5 |
| HIP1 | Vic11 | 1_4_1 | IL18 | 20 |
| HIP1 | Vic11 | 1_0_1 | IL18 | 16.5 |
| HIP1 | Vic11 | 0_4_1 | IL18 | 28.75 |
| HIP1 | Vic11 | 0_0_1 | IL18 | 18 |
| HIP9 | Mock | 1_4_1 | IL18 | 24 |
| HIP9 | Mock | 1_0_1 | IL18 | 16 |
| HIP9 | Mock | 0_4_1 | IL18 | 16.5 |
| HIP9 | Mock | 0_0_1 | IL18 | 22 |
| HIP9 | Cal09 | 1_4_1 | IL18 | 19 |
| HIP9 | Cal09 | 1_0_1 | IL18 | 18 |
| HIP9 | Cal09 | 0_4_1 | IL18 | 28.5 |
| HIP9 | Cal09 | 0_0_1 | IL18 | 20.5 |
| HIP9 | Vic11 | 1_4_1 | IL18 | 17.25 |
| HIP9 | Vic11 | 1_0_1 | IL18 | 16.5 |
| HIP9 | Vic11 | 0_4_1 | IL18 | 35.5 |
| HIP9 | Vic11 | 0_0_1 | IL18 | 16.5 |
| HIP18 | Mock | 1_4_1 | IL18 | 27 |
| HIP18 | Mock | 1_0_1 | IL18 | 22.75 |
| HIP18 | Mock | 0_4_1 | IL18 | 30.5 |
| HIP18 | Mock | 0_0_1 | IL18 | 15 |
| HIP18 | Cal09 | 1_4_1 | IL18 | 45.5 |
| HIP18 | Cal09 | 1_0_1 | IL18 | 27.5 |
| HIP18 | Cal09 | 0_4_1 | IL18 | 16 |
| HIP18 | Cal09 | 0_0_1 | IL18 | 15.5 |
| HIP18 | Vic11 | 1_4_1 | IL18 | 19.5 |
| HIP18 | Vic11 | 1_0_1 | IL18 | 19.5 |
| HIP18 | Vic11 | 0_4_1 | IL18 | 29.25 |
| HIP18 | Vic11 | 0_0_1 | IL18 | 16.5 |
| HIP1 | Mock | 1_4_1 | TNF | 3074.5 |
| HIP1 | Mock | 1_0_1 | TNF | 322 |
| HIP1 | Mock | 0_4_1 | TNF | 236 |
| HIP1 | Mock | 0_0_1 | TNF | 27.5 |
| HIP1 | Cal09 | 1_4_1 | TNF | 381.75 |
| HIP1 | Cal09 | 1_0_1 | TNF | 908 |
| HIP1 | Cal09 | 0_4_1 | TNF | 77.5 |
| HIP1 | Cal09 | 0_0_1 | TNF | 20 |
| HIP1 | Vic11 | 1_4_1 | TNF | 196.75 |
| HIP1 | Vic11 | 1_0_1 | TNF | 316.5 |
| HIP1 | Vic11 | 0_4_1 | TNF | 201.5 |
| HIP1 | Vic11 | 0_0_1 | TNF | 22 |
| HIP9 | Mock | 1_4_1 | TNF | 4424 |
| HIP9 | Mock | 1_0_1 | TNF | 33.5 |
| HIP9 | Mock | 0_4_1 | TNF | 50.75 |
| HIP9 | Mock | 0_0_1 | TNF | 33.25 |
| HIP9 | Cal09 | 1_4_1 | TNF | 191.5 |
| HIP9 | Cal09 | 1_0_1 | TNF | 88 |
| HIP9 | Cal09 | 0_4_1 | TNF | 124.25 |
| HIP9 | Cal09 | 0_0_1 | TNF | 41.75 |
| HIP9 | Vic11 | 1_4_1 | TNF | 154.5 |
| HIP9 | Vic11 | 1_0_1 | TNF | 73 |
| HIP9 | Vic11 | 0_4_1 | TNF | 66 |
| HIP9 | Vic11 | 0_0_1 | TNF | 18.25 |
| HIP18 | Mock | 1_4_1 | TNF | 2118 |
| HIP18 | Mock | 1_0_1 | TNF | 2171.5 |
| HIP18 | Mock | 0_4_1 | TNF | 256.25 |
| HIP18 | Mock | 0_0_1 | TNF | 17 |
| HIP18 | Cal09 | 1_4_1 | TNF | 327.25 |
| HIP18 | Cal09 | 1_0_1 | TNF | 427.5 |
| HIP18 | Cal09 | 0_4_1 | TNF | 116 |
| HIP18 | Cal09 | 0_0_1 | TNF | 15.5 |
| HIP18 | Vic11 | 1_4_1 | TNF | 168.25 |
| HIP18 | Vic11 | 1_0_1 | TNF | 238.75 |
| HIP18 | Vic11 | 0_4_1 | TNF | 103.75 |
| HIP18 | Vic11 | 0_0_1 | TNF | 19.5 |
| HIP1 | Mock | 1_4_1 | IL21 | 28 |
| HIP1 | Mock | 1_0_1 | IL21 | 33 |
| HIP1 | Mock | 0_4_1 | IL21 | 30 |
| HIP1 | Mock | 0_0_1 | IL21 | 34.5 |
| HIP1 | Cal09 | 1_4_1 | IL21 | 48 |
| HIP1 | Cal09 | 1_0_1 | IL21 | 46.5 |
| HIP1 | Cal09 | 0_4_1 | IL21 | 56.5 |
| HIP1 | Cal09 | 0_0_1 | IL21 | 48.5 |
| HIP1 | Vic11 | 1_4_1 | IL21 | 35.25 |
| HIP1 | Vic11 | 1_0_1 | IL21 | 31 |
| HIP1 | Vic11 | 0_4_1 | IL21 | 25 |
| HIP1 | Vic11 | 0_0_1 | IL21 | 31 |
| HIP9 | Mock | 1_4_1 | IL21 | 29.5 |
| HIP9 | Mock | 1_0_1 | IL21 | 35.5 |
| HIP9 | Mock | 0_4_1 | IL21 | 29 |
| HIP9 | Mock | 0_0_1 | IL21 | 36.5 |
| HIP9 | Cal09 | 1_4_1 | IL21 | 26.75 |
| HIP9 | Cal09 | 1_0_1 | IL21 | 29 |
| HIP9 | Cal09 | 0_4_1 | IL21 | 26.5 |
| HIP9 | Cal09 | 0_0_1 | IL21 | 25 |
| HIP9 | Vic11 | 1_4_1 | IL21 | 30.5 |
| HIP9 | Vic11 | 1_0_1 | IL21 | 40.5 |
| HIP9 | Vic11 | 0_4_1 | IL21 | 43.5 |
| HIP9 | Vic11 | 0_0_1 | IL21 | 40.5 |
| HIP18 | Mock | 1_4_1 | IL21 | 30 |
| HIP18 | Mock | 1_0_1 | IL21 | 29.5 |
| HIP18 | Mock | 0_4_1 | IL21 | 28.25 |
| HIP18 | Mock | 0_0_1 | IL21 | 29.5 |
| HIP18 | Cal09 | 1_4_1 | IL21 | 28.5 |
| HIP18 | Cal09 | 1_0_1 | IL21 | 34.75 |
| HIP18 | Cal09 | 0_4_1 | IL21 | 30 |
| HIP18 | Cal09 | 0_0_1 | IL21 | 35.5 |
| HIP18 | Vic11 | 1_4_1 | IL21 | 23.75 |
| HIP18 | Vic11 | 1_0_1 | IL21 | 41.5 |
| HIP18 | Vic11 | 0_4_1 | IL21 | 25 |
| HIP18 | Vic11 | 0_0_1 | IL21 | 26.5 |
